# Specialized pathogenic cells release Tc toxins using a type 10 secretion system

**DOI:** 10.1101/2023.02.22.529496

**Authors:** Oleg Sitsel, Zhexin Wang, Petra Janning, Lara Kroczek, Thorsten Wagner, Stefan Raunser

## Abstract

Disease-causing bacteria use a variety of secreted toxins to invade and subjugate their hosts. While the machinery responsible for secretion of many smaller toxins has already been established, it remains enigmatic for larger ones such as Tc toxins from human and insect pathogens, which approach the size of a prokaryotic ribosome. In the present study, we combine targeted genomic editing, proteomic profiling and cryo-electron tomography of the insect pathogen *Yersinia entomophaga* to reveal that a specialized subset of bacterial cells produces the Tc toxin YenTc as part of a complex toxin cocktail released into the environment by controlled cell lysis using a transcriptionally-coupled, pH-dependent type 10 secretion system (T10SS). Our results dissect the process of Tc toxin export by a T10SS in hitherto unprecedented detail, identifying that T10SSs operate via a previously unknown lytic mode of action, and establishing them as crucial players in the size-insensitive release of cytoplasmically folded toxins. With T10SSs directly embedded in Tc toxin operons of major human pathogens such as *Yersinia pestis* and *Salmonella enterica*, we anticipate our findings to model an important aspect of pathogenesis in bacteria with a significant impact on global human health.

## Introduction

When attacking eukaryotes, pathogenic bacteria secrete an array of toxic proteins that modulate the defences of the host on a cellular level allowing the pathogen to establish an infection. Tc toxins are a megadalton-scale, architecturally complex (*1*) example of such proteins, for which the secretion mechanism remains highly enigmatic (*2*). Originally discovered in the insecticidal bacterium *Photorhabdus luminescens* (*3*), these non-phage-derived protein complexes are now known to be crucial virulence factors in numerous highly prominent insect and human pathogens (*4*). The structure of Tc toxins and the roles of their three core subunits TcA, TcB and TcC have been extensively investigated in the past several years (*1, 5-10*). While the details of Tc complex assembly and secretion in a biological context are so far missing (*1*), evidence suggests that secreted Tc toxins first attach to their target cells via glycosylated proteinaceous receptors (*11*) and other surface exposed glycan moieties (*10, 12-14*) and then undergo subsequent endocytosis. Endosomal acidification triggers a conformational change by collapsing the linker domains of the five TcA protomers, which drives the central TcA channel into the endocytic membrane (*8*). The membrane-embedded tip of the channel then opens (*7*), releasing a cytotoxic moiety originally stored in the TcB-TcC cocoon into the target cell cytoplasm (*1*), where it modifies substrates such as Rho GTPases and the actin cytoskeleton, ultimately triggering cell death (*1, 15, 16*).

As with other toxins of Gram-negative bacteria, Tc toxins face the challenge of crossing three major barriers when exiting the cell: the phospholipid inner membrane, peptidoglycan sacculus, and lipopolysaccharide outer membrane (*17*). Prior experiments have led to the existence of several competing hypotheses regarding Tc toxin secretion. TcB-/TcC-driven translocation of a *P. luminescens* Tc toxin via an auto-transport mechanism (*18, 19*) with attachment to the bacterial cell surface (*20*) and subsequent lipase-assisted release into the surrounding environment (*18*) has been proposed on one hand. On the other, secretion of the *Yersinia pestis* and *Yersinia entomophaga* Tc toxins has been suggested to occur via the T3SS and outer membrane vesicles (OMVs), respectively (*21, 22*). For other toxins in Gram-negative bacteria, it has so far been established that they use the type 1 secretion system (T1SS), Tat/T2SS, Sec/T2SS, Sec/T5SS and outer membrane vesicle (OMV) pathways to traverse into extracellular space, or the T3SS, T4SS and T6SS pathways to inject directly into host target cells (*17, 23*), with use of each secretion system introducing unique requirements to its substrates such as cytoplasmic unfolding and presence of a specific signal sequence (*17*). The physical dimensions of these secretion systems also limits the vast majority of toxins to a size of tens of kilodaltons (kDa), although certain larger cargos such as the 900 kDa RTX toxins can still be threaded through a T1SS, a feat afforded by their cytoplasmically unfolded repeat domain architecture (*24*). Alternative exit routes from the cell also exist. Shiga toxins are for instance directly encoded on fully functional bacteriophage genomes (which also limits their size), and they make use of timed lytic viral release to escape from bacterial cells (*25*). This process is mediated by holin/endolysin/spanin-containing phage lysis cassettes (*26*): upon reaching a critical concentration, phage holins spontaneously form oligomeric pores in the inner membrane that either translocate a cytoplasmic endolysin to the periplasm or stimulate the refolding of a periplasmic membrane-bound endolysin into an active state. This then degrades the peptidoglycan layer, allowing the spanin complex to fuse the inner and outer membranes and thereby enable viral escape (*27*). Holin/endolysin/spanin clusters reminiscent of classical phage lysis cassettes but outside of a bacteriophage genomic context were very recently established as the so-called type 10 secretion system (T10SS) (*28*). They have been proposed to mediate protein export through a non-lytic mode of action (*28-30*), with the archetypal *Serratia marcescens* T10SS being responsible for secretion of chitinolytic machinery used to survive on chitin as a sole carbon source (*30, 31*).

In this work, we aimed to determine which of these numerous possibilities Tc toxins use for secretion. For this we focused on *Y. entomophaga*, a relative of the deadly human pathogen *Y. pestis* that has instead evolved to target insects. The virulence factor that makes *Y. entomophaga* extremely lethal to insects is its sole Tc toxin, YenTc, which it secretes into the environment (*22*). The intricate multi-subunit structure of this toxin (*10*) gives rise to a complex of roughly 2.4 MDa, which is comparable in size to the 2.5 MDa prokaryotic ribosome. The extraordinary size and complexity of YenTc represent significant challenges for the secretion machinery established for other much smaller toxins. We used a cutting-edge combination of microscopy, proteomic analyses and targeted genomic knockouts to reveal that a small subset of specialized cells, which we term soldier cells, releases YenTc as well as numerous other virulence factors and toxins using a pH-sensitive T10SS in a lytic fashion. We identified the regulatory machinery that controls the soldier cell phenotype in a temperature-sensitive manner and couples production of YenTc and other virulence factors to the T10SS. Using cryo-electron tomography (cryo-ET) we visualized the step-by-step release of YenTc. These results resolve the mystery surrounding Tc toxin secretion in insect and human pathogens, and demonstrate how specialized toxin-producing cells confer virulence to an entire bacterial population.

## Results

### pH-dependent secretion of Tc toxins

In order to investigate the secretion of Tc toxins, we started by exploring the influence of growth media composition on YenTc production and secretion. We immediately noticed that *Y. entomophaga* secretes very poorly into media that acidify during cell growth compared to non-acidifying media (**Fig. S1a**). This led us to hypothesize that protein secretion from *Y. entomophaga* is pH-dependent, and indeed raising the pH of the acidified medium from 5.5-6.0 to 7.0-8.0 or reconstituting the cells in a buffer with elevated pH caused them to secrete a multitude of proteins within 10 minutes (**Fig. S2a**). We systematically screened the pH dependence of secretion and found that it occurs in the range of pH 6.3-10.0, with an optimum between pH 6.9-8.9 (**Fig. S2b-c**). Many insect families, including Coleoptera to which the natural *Y. entomophaga* host *Costelytra zealandica* belongs, have an acidic anterior midgut and a more alkaline posterior midgut (*32, 33*). The identified pH sensitivity of secretion would cause release of YenTc near the latter, which is where invading *Y. entomophaga* were found to actively establish themselves (*34*).

We next turned to mass spectrometry to identify if other proteins are co-secreted with YenTc. Surprisingly, the secreted fraction contained numerous proteins that are not typically found in an extracellular context (**Fig. 1a, inset**), while a further comparison of proteins in the secreted fraction to those remaining in the cells revealed that the former was highly enriched not only in YenTc, but also in a variety of other toxins and virulence factors such as PirAB, NucA and chitin-modifying enzymes (**Fig. 1a, Source Data file**). Mysteriously, nearly all lack an established signal sequence for export (**Fig. S3a**). Taken together, this indicates that *Y. entomophaga* produces a protein cocktail which is exported by a protein secretion machinery that does not require its cargoes to possess a dedicated secretion signal sequence.

**Figure 1.**
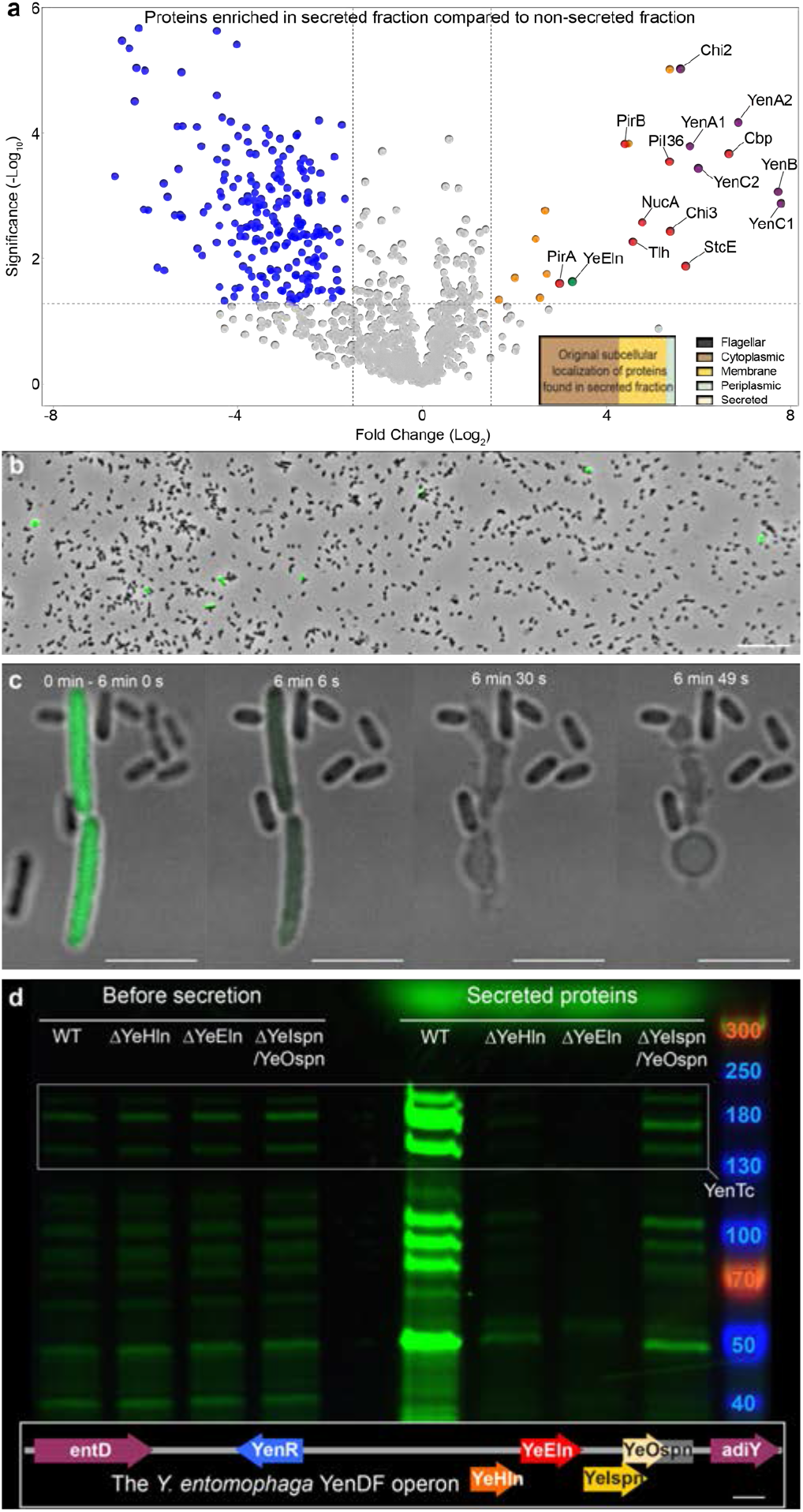
*Y. entomophaga* soldier cells release YenTc and other toxins using YenDF, a T10SS. **a**, Volcano plot of the MS analysis showing significant enrichment (t-test p-value < 0.05) of toxic proteins and a T10SS component in the secreted protein fraction compared to the non-secreted fraction. Purple: YenTc components. Red: other secreted toxins and virulence factors. Green: YenDF structural components. Orange: cytoplasmic proteins enriched in the secreted fraction. The relevant UniProt accession numbers are provided in the Materials and methods section. n = 3 biological replicates. The full proteomic datasets used to generate this figure are available in the Source Data file. Inset: the original subcellular localizations of those proteins found in the secreted fraction for which such information is available (540 of 1656 total hits). **b**, Only a minor fraction of an isogenic *Y. entomophaga* population produces YenTc-sfGFP (green). Scale bar: 20 μm. **c**, Timelapse of YenTc-sfGFP release from soldier cells (green) after the pH of acidified growth media was raised to 8.0. Scale bars: 5 μm. **d**, Knockout of YenDF components blocks secretion of *Y. entomophaga* soldier cells upon raising the pH of acidified growth media. The bands corresponding to YenA1, YenA2 and YenB are boxed for clarity. Inset: structure of the YenDF operon. Scale bar: 200 bp.

### Soldier cells secrete YenTc using a T10SS

To identify the pathway these bacteria employ to release YenTc and possibly other constituents of the toxic cocktail, we established a targeted genomic editing protocol for *Y. entomophaga* that can be used to make scarless edits (**Fig. S1b**). After knocking out all established secretion systems in *Y. entomophaga* (Sec, Tat, T1SS, T2SS, T3SS, T6SS), as well as components of the non-standard secretion pathway hypothesized for the *P. luminescens* Tc toxin (*18, 19*) (lipases, OMV-promoting enzymes, the Tc toxin itself), all resulting strains secreted YenTc and/or other toxins (**Fig. S4**), meaning that neither of the previously proposed mechanisms nor other common secretion systems are used for export of these toxins from *Y. entomophaga*.

To resolve this mystery, we proceeded by fusing the YenA1 component of YenTc with sfGFP in order to directly monitor secretion of the toxin. Fluorescence microscopy revealed the surprising fact that only a small subpopulation of these isogenic cells expressed the toxin at any given time (**Fig. 1b**), indicating bimodality (*35*). Furthermore, the toxin-expressing cells were often morphologically distinct from the rest in the post-log phase (**Fig. 1c**), being larger and noticeably less motile. Of all factors tested (quorum sensing, oxygen levels, genotoxic stress, temperature etc.), lower temperature had by far the strongest influence on the propensity of cells to become YenTc producers (**Fig. S5a-n**), in line with the observation that *Y. entomophaga* does not secrete this toxin into growth media at temperatures above 30 °C (*22*). This makes biological sense for such an insect pathogen, since any organism it may encounter with a highly elevated body temperature is unlikely to be a suitable host. Absence of another host marker, complex organic compounds, also strongly disincentivized differentiation into YenTc-producing cells (**Fig. S5h, n**). Finally, we reduced the number of YenTc-expressing cells in the population by two thirds when we knocked out production of autoinducer-1 quorum sensing molecules. This showcases the importance of interbacterial communication in the decision to produce YenTc, and interestingly correlates with the reduced secretion of a corresponding hit from a transposon mutagenesis assay (*36*).

We then used confocal fluorescence microscopy to visualize the pH-induced secretion of YenTc-expressing cells. While cells not expressing YenTc remained unaltered, the enlarged rod-like morphology of YenTc-expressing cells underwent a striking metamorphosis in a matter of minutes, collapsing into an associated cluster of vesicles and releasing the previously cytoplasmically localized YenTc in the process (**Fig. 1c**). Thus, the cells that produce YenTc are sacrificed for the release of the toxin. This demonstrates that the pH-dependent secretion of YenTc is actually not a typical secretion process, but the result of a controlled lysis strictly dedicated to toxin release. This remarkable functional and morphological differentiation of YenTc-producing cells compared to their naive counterparts led us to term them soldier cells. In the light of soldier cells using controlled lysis to release YenTc, we found the exclusive appearance of an M15 family metalloprotease with predicted endolysin activity in the secreted proteome to be particularly intriguing (**Fig. 1a, Source Data file**) due to its high similarity to the corresponding component of the archetypal *S. marcescens* T10SS (*28, 29*). Analyzing the *Y. entomophaga* genome, we found that similarly to the *S. marcescens* T10SS, the metalloprotease is encoded between a holin and bicomponent spanin (**Fig. 1d, inset**) flanked by transcription factors and various housekeeping genes, and thus indeed belongs to a T10SS discrete from the YenTc pathogenicity island. Studies of its *Serratia marcescens* counterpart indicate (*29, 37*) that the metalloprotease acts as an L-alanyl D-glutamyl endopeptidase that cleaves peptidoglycan cross-links and therefore indeed represents an endolysin. We termed the endolysin YeEln, the holin YeHln, and the spanins YeIspn and YeOspn, respectively. Our suspicion that we were on the right track was strengthened by the results of an independent *Y. entomophaga* transposon mutagenesis assay (*36*), where some hits with reduced protein secretion were associated with an intergenic region close to the gene encoding YeHln. Remarkably, we completely eliminated *Y. entomophaga* protein export when we deleted the T10SS fully (**Fig. S4**) as well as its YeHln and YeEln components individually (**Fig. 1d**). We also found that YeIspn and YeOspn knockouts are largely but not completely defective when we applied shearing forces to the frail cells generated by YeEln activity (**Fig. S8**), and we later show that these spanins are crucial in more natural contexts where shearing forces are not involved (*38*) (**Fig. 3**). These results demonstrate that this T10SS is responsible for the release of not only YenTc by controlled lysis but also of all other toxic proteins we found enriched in the secreted fraction, explaining why nearly all lack a dedicated secretion signal sequence (**Fig. S3a**) and prompting us to name it the *Yersinia entomophaga* death factor (YenDF).

### Synchronization of YenDF and YenTc production

We next aimed to determine how soldier cells activate production of YenDF, with an OmpR/PhoB-type helix-turn-helix fold protein that we termed YenR encoded directly upstream of YeHln drawing our attention as a potential candidate for controlling YenDF expression, analogously to how the LysR-type transcription factor ChiR controls bimodal production of the nearby *S. marcescens* T10SS and several chitinolytic enzymes (*30*). To test this, we replaced its native promoter with arabinose-inducible regulatory elements to create the Ara-YenR strain. In the absence of arabinose, this strain completely stopped secreting and – surprisingly – producing YenTc (**Fig. 2a**). Adding arabinose to Ara-YenR massively boosted both YenTc production and protein secretion compared to wild type cells, while the control Ara-YenR ΔYenDF strain generated very high levels of intracellular YenTc that it was unable to secrete (**Fig. 2a**).

**Figure 2.**
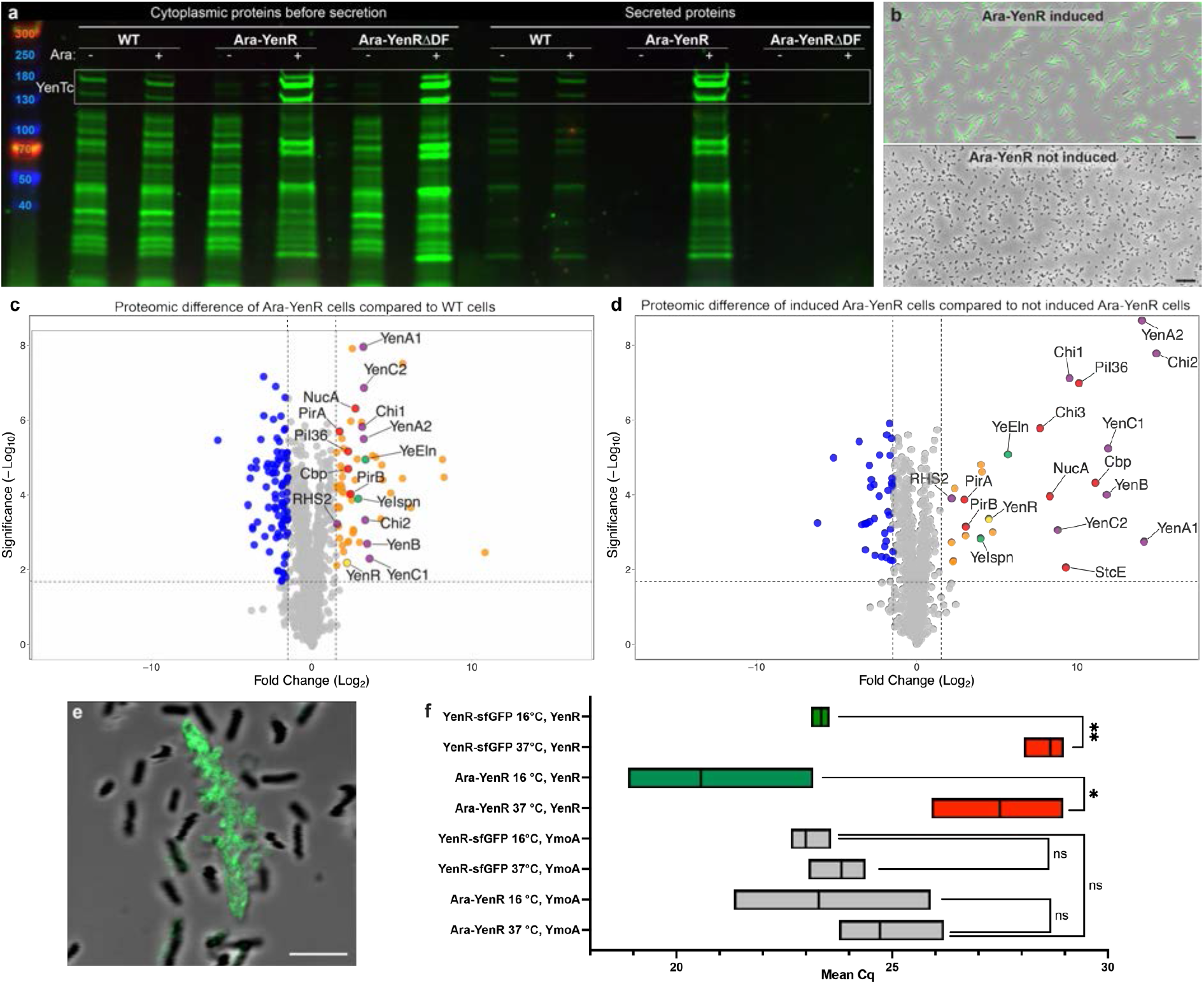
Soldier cells use YenR to synchronize the expression of YenDF with that of a deadly cocktail of toxins in a temperature-sensitive manner. **a**, Arabinose induction of the Ara-YenR strain causes a massive increase in toxin production and secretion compared to wild type (WT) cells, while absence of induction abolishes these completely. **b**, Induction of YenR in an Ara-YenR YenTc-sfGFP strain causes all cells to convert into a soldier cell phenotype (green), while in absence of induction no soldier cells appear. Scale bar: 20 μm. **c-d**, Volcano plot showing significant enrichment (t-test p-value < 0.02) of toxic proteins and YenDF components in the total protein fraction of induced Ara-YenR cells compared to WT cells (c) or non-induced Ara-YenR cells (d). Colors match Fig. 1a, additionally the regulatory protein YenR is marked yellow. n = 3 biological replicates each. The full proteomic datasets used to generate these figures are available in the Source Data file. **e**, Only explosion-competent soldier cells of the YenR-sfGFP strain express sufficient YenR-sfGFP to be seen by confocal fluorescence microscopy. Scale bar: 5 μm. **f**, YenR transcript levels as measured by RT-qPCR are strongly dependent on temperature regardless of whether the 5’ or 3’ UTR is disrupted (in the Ara-YenR and YenR-sfGFP strains, respectively) while mRNA levels of the YmoA control are not. Data is shown as mean ± minimal/maximal values, n = 3 biological replicates, with p-values measured by t-test.

To validate whether YenR indeed controls both YenDF and YenTc production as these phenotypes suggest, we analysed the cytoplasmic contents of induced Ara-YenR cells by mass spectrometry. Remarkably, our comparison with wild type and non-induced Ara-YenR cells (**Fig. 2c-d, Source Data file**) revealed that YenR controls production of not only YenDF components and YenTc, but also the numerous other secreted toxins and virulence factors we identified earlier (**Fig. 1a**). This demonstrates that YenTc-expressing soldier cells, rather than yet another specialized cell subpopulation, are the source of this lethal protein cocktail. Promoter region analysis of these YenR-controlled genes indicates the presence of an overrepresented sequence that may function as the YenR binding site (**Fig. S3b**).

We then imaged an induced Ara-YenR YenTc-sfGFP strain, and found that the entire bacterial population converted into secretion-competent soldier cells (**Fig. 2b, Movie S1**). This means that YenR is the key to both coupling production of YenTc to the production of its secretion system and generating the soldier cell phenotype. To investigate whether the appearance of soldier cells is induced by high intracellular levels of YenR also in a native context, we monitored populations of a YenR-sfGFP strain by confocal fluorescence microscopy. Indeed, only the large, explosion-capable soldier cells showed an appreciable level of YenR-sfGFP fluorescence (**Fig. 2e**). Unlike the soluble YenTc-sfGFP toxin (**Fig. 1c**), the DNA-bound YenR-sfGFP transcription factor remained largely bound to the cell remnants after YenDF-mediated lysis.

Interestingly, induction of the Ara-YenR strain at a growth temperature of 37 °C instead of 20 °C did not generate soldier cells (**Fig. S2d-h**). This is in line with our previous observation that soldier cells are absent in cultures grown at elevated temperatures (**Fig. S5m, n**), indicating that a layer of thermosensitive regulation is at play that is possibly reminiscent to that of the *Y. enterocolitica* Tc toxin LysR-type transcriptional regulator TcaR2, which is regulated at the protein level by temperature-sensitive degradation (*39*). Based on the observation that only cells with increased YenR levels differentiate (**Fig. 2e**), we hypothesized that elevated temperatures affect YenR production either at the mRNA or protein level. To discriminate between these possibilities, we undertook RT-qPCR analyses on total RNA extracted from the induced Ara-YenR strain (with a disrupted native YenR 5’ UTR) and YenR-sfGFP strain (with a disrupted native YenR 3’ UTR) grown at either 16 °C or 37 °C. YenR transcript levels at 37 °C dropped significantly compared to 16 °C in both strains as opposed to the non-affected control gene (**Fig. 2f**). This suggests the presence of a heat-repressible RNA thermostat in the coding sequence of YenR that affects mRNA stability similarly to what has been described for the cold-inducible CspA mRNA in *Escherichia coli* (*40*). Thermosensitivity of *Y. entomophaga* differentiation is thus controlled already at the YenR transcript level, explaining how the soldier cell phenotype is suppressed in conditions incompatible with insect hosts.

### The YenTc release mechanism

We next investigated if the holin YeHln is needed for the endolysin YeEln to reach the periplasm and which role the pH trigger plays in this process. For this we monitored YeEln mediated, pH-dependent protein release in Ara-YenR cells with or without intact YeHln. Only cells containing both YeHln and YeEln were able to release proteins upon elevation of pH (**Fig. S6a**), suggesting that YeEln uses YeHln to reach the periplasm as seen in the holin/endolysin/spanin systems of bacteriophages (*26*). A logical conclusion would be that elevation of extracellular pH triggers the activation of holin transport in soldier cells. This is in contrast to the situation in phage infections, where holin levels in the bacterial inner membrane increase until reaching a critical allele-timed concentration at which a local decrease in the proton motive force (PMF) spontaneously triggers them to form large pores (*41-43*). However, increasing external pH can also decrease the PMF between the cytoplasm and periplasm (*44*), so that the triggering of YeHln by elevation of external pH could also be explained by an accompanying decrease in the PMF (*44*). This interpretation is also supported by our observation of controlled lysis in Ara-YenR cells under sudden anaerobiotic stress (**Fig. S6b**), which causes a loss in PMF due to decrease of cytoplasmic pH (*45*). This means that although the initial triggers are different, the ultimate cause of holin activation is likely to be similar in phage holins and YenDF.

In order to visualize the individual steps of YenDF-mediated YenTc release from soldier cells following pH triggering, we produced derivatives of the Ara-YenR strain blocked in the pre-secretion state as well as the post-holin, post-endolysin and post-spanin states by knockout of the corresponding YenDF components. We then vitrified the cells by plunge-freezing into liquid ethane. Given the thickness of the cells from the first three strains (**Fig. S7a**), we then prepared 50-100 nm thin lamellae ideally suited for cryo-ET using cryo-focused ion beam (cryo-FIB) milling (*46*) (**Fig. S7c-f**), while the cells in the post-spanin state could be imaged directly. It is important to emphasize that the 10-15 kDa YenDF components themselves are far too small to be visualized *in situ*, therefore we instead aimed to analyze the effects of each component’s action on cellular ultrastructure. The tomograms revealed that the cell envelope is fully intact in the pre-secretion state (**Fig. 3a and e, Movie S2**). Strikingly, we found many fully assembled YenTc holotoxins composed of all three subunits (TcA, TcB, and TcC) in the cytoplasm (**Fig. 3a, inset**) which finally establishes the bacterial cytoplasm as the location of holotoxin assembly for Tc toxins.

**Figure 3.**
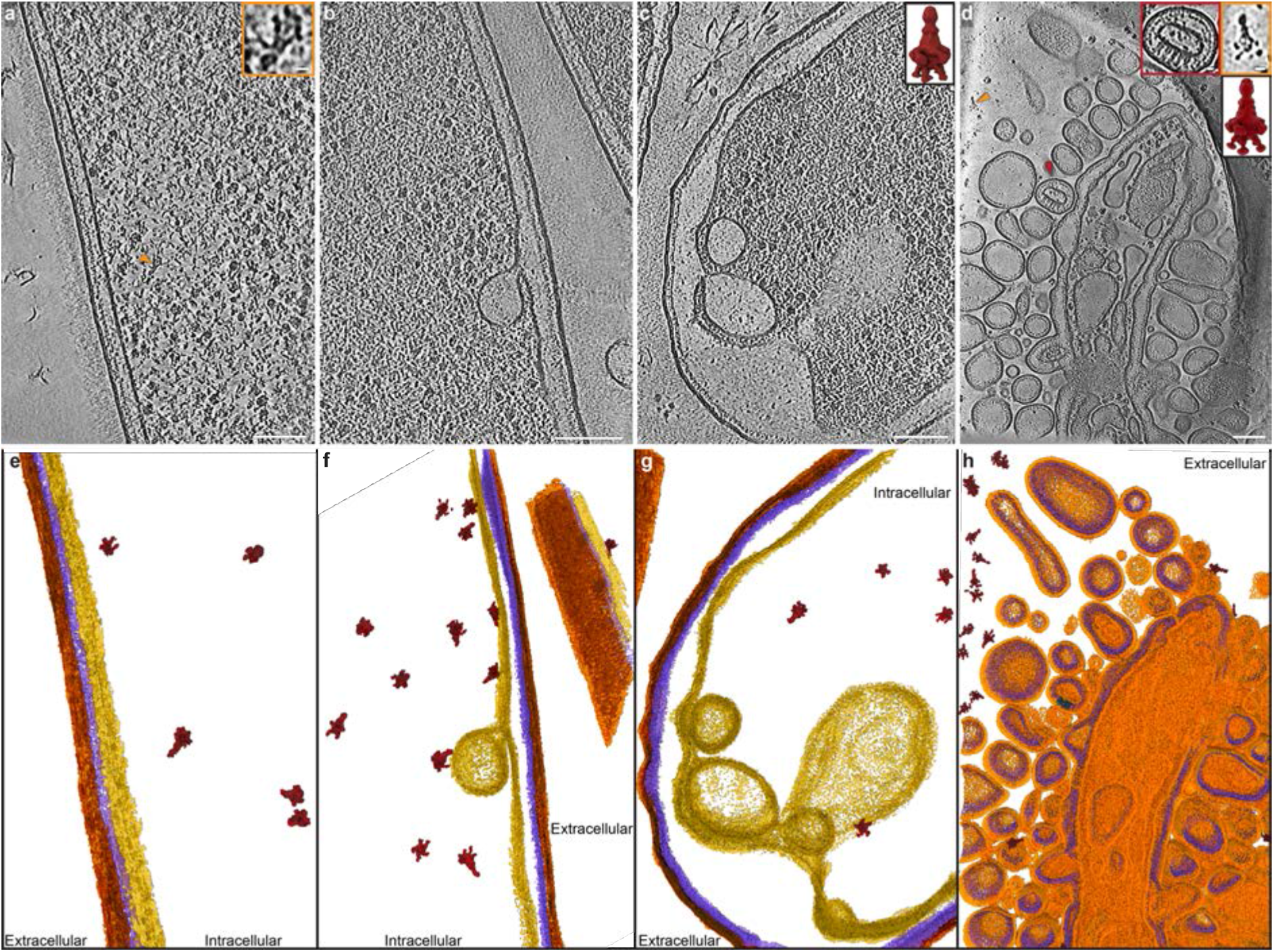
Step-by-step effect of YenDF action and release of YenTc visualized by cryo-ET. **a**, A single slice from an Ara-YenR ΔYenDF cell tomogram, representing the pre-secretion state. Inset: a fully assembled cytoplasmic YenTc holotoxin. An orange arrowhead denotes its position in the tomogram slice. Scale bar: 10 nm. **b**, Ara-YenR ΔYenEln/YeIspn/YeOspn cell tomogram slice, representing the state of secretion after holin action. **c**, Ara-YenR ΔYeIspn/YeOspn cell tomogram slice, representing the state of secretion after endolysin action. Inset: Structure of the intracellular YenTc holotoxin complex from Ara-YenR ΔYeIspn/YeOspn cells, determined by subtomogram averaging after automated particle picking by TomoTwin. See Figs. S9-S10 for more details. **d**, Ara-YenR cell tomogram slice, representing the state of secretion after spanin action. Left inset: unfused vesicle derived from an area of the cell envelope that contained proteins spanning the entire cell wall. A red arrowhead denotes its position in the tomogram slice. Scale bar: 10 nm. Right inset: A YenTc holotoxin secreted to the external environment. An orange arrowhead denotes its position in the tomogram slice. Scale bar: 10 nm. Lower inset: Structure of the secreted YenTc holotoxin complex determined by subtomogram averaging after manual picking. See Fig. S10 for more details. Scale bars in all tomogram slice views: 100 nm. **e-g**, Annotated densities from the tomograms shown in (**a-d**). Dark orange: outer membrane, purple: peptidoglycan, yellow: inner membrane, dark red: YenTc. **h**, Annotated densities from the tomogram shown in (**d**), presented as a diagonally sectioned view to demonstrate the internal structure that arises after spanin action. Light orange: fused membranes, purple: peptidoglycan, dark red: YenTc, dark blue: cell envelope-spanning protein complexes.

After the pH trigger, the cells in the post-holin state demonstrate that YeHln action leads to perturbations of the inner membrane in the form of invaginations not seen in ΔYeHln cells (**Fig. 3b and f, Movie S3, Fig. S7b**). However, we did not observe micron-scale lesions with accompanying expulsion of cytoplasmic material into the periplasm that have been described for the activated form of archetypal phage holin S105 (*41*), even when intact cells were examined to exclude the possibility that cryo-FIB milling had removed areas containing such features (**Fig. S7a and b**). To validate the presence of YeHln, we used confocal fluorescence microscopy to determine the distribution of its fluorescently tagged variant and found that YeHln distributes throughout the cell surface in nearly ubiquitous small foci (**Fig. S6c**) reminiscent of the oligomeric “rafts” described for S105 (*43, 47*). A very recently proposed alternative mode of holin action does not require pore formation but rather utilizes a membrane weakening/flipping mechanism for endolysin transport (*48*). Based on the absence of observed lesions here, YeHln may represent an example of such a holin, making *Y. entomophaga* an excellent future testbed for investigating this exciting hypothesis.

Next, we analyzed the tomograms of the cells in the post-endolysin state to understand the cellular effects of the enzymatic action of YeEln. The inner membrane of these cells is largely detached from the peptidoglycan layer and outer membrane, resulting in severe inward bending of the inner membrane (**Fig. S8a, Fig. 3c and g, Movie S4**). In a spanin-free situation, the action of YeEln eventually leads to such significant weakening of the peptidoglycan sacculus that internal osmotic pressure (*49*) drives the cell to expand into a spheroplast (**Fig. S8b-c**). Our tomograms show that these spheroplasts have significantly lower protein density compared to pre-spheroplast cells due to their larger volume, making them a highly attractive target for future *in situ* structural proteomics studies (**Fig. S8d, Figs. S9-S10**).

When the spanins YenIspn and YenOspn are however present, the bacterium undergoes a dramatic transformation that converts it into a loosely bound cluster of membrane vesicles (**Fig. 3d and h, Movie S5**). Membranes in areas with cell envelope-spanning protein complexes stay discrete after this secretion step (**Fig. 3d, left inset**) and the cell wall of the vesicles appears intact, indicating that this process does not disrupt the lipoprotein Lpp tether of peptidoglycan to the former outer membrane (**Fig. 3d**). Importantly, this spanin-driven metamorphosis releases YenTc holotoxins (**Fig. 3d, right and bottom insets, Fig. S10**), indicating that the spanins not only trigger the fusion of the outer and inner membrane of soldier cells, leading to formation of unimembrane vesicles, but that this process also releases the cytoplasmic contents into the external environment.

### T10SSs operate via a hitherto undescribed lytic mechanism

The novel lytic mechanism of T10SS-mediated YenTc release we identified is at odds with the non-lytic mode of action proposed for the archetypal T10SS from *Serratia marcescens* (*28-30*). Although the primary focus of this study is on the identification and characterization of the Tc toxin secretion mechanism, we were intrigued to determine whether the modes of *Y. entomophaga* and *S. marcescens* T10SS action are indeed so different. To do so, we applied the targeted genomic editing protocol we originally developed for *Y. entomophaga* (**Fig. S1b**) to the well-characterized *S. marcescens* type strain in order to create an arabinose-inducible Ara-ChiR strain functionally analogous to the *Y. entomophaga* Ara-YenR strain. The secretion of the Ara-ChiR strain had two important differences compared to that of *Y. entomophaga* Ara-YenR grown in the same conditions: it was no longer inhibited by acidification of the media and its temperature sensitivity profile was inverted (**Fig. S11a**), showcasing how differences in the lifestyle of these two bacterial pathogens impose variation on the temperature of T10SS/cargo production and pH sensitivity of secretion. As in the case of *Y. entomophaga* (**Fig. 1a, inset**), the *S. marcescens* Ara-ChiR secreted fraction was full of typically non-exported proteins (**Fig. S12a inset, Fig. S12c, Source Data file**) in addition to the abundantly present chitinolytic machinery, and release of these proteins was dependent on the *S. marcescens* T10SS, as evidenced by their absence in the secreted fraction of the corresponding T10SS knockout strain (**Fig. S11a, Fig. S12b-c, Source Data file**). As we found that we could controllably trigger the T10SS by anaerobiotic stress-induced proton motive force collapse (**Fig. S11c, Movie S6**) similarly to *Y. entomophaga* soldier cells (**Fig. S6b, Movie S1**), we again acquired cryo-electron tomograms directly on vitrified samples in order to visualize the triggered Ara-ChiR cells in much greater detail (**Fig. S11d**). The tomograms revealed that they undergo the same dramatic spanin-driven transformation as *Y. entomophaga* soldier cells (**Fig. S11e, Movies S7-S8**) with the bacteria converting into unimembrane vesicles, forcibly expelling their cytoplasmic contents and thus allowing rapid release of accumulated chitinolytic machinery. This establishes that T10SSs mediate lytic protein release rather than operate in the non-lytic fashion that was previously proposed (*29*), providing a much-needed mechanistic explanation to the previously unresolved issue of how T10SS-exported proteins cross the outer membrane after protein maturation (*28*) and finally revealing the biological rationale of why only a minor fraction of *S. marcescens* cells in a population express the T10SS (*29, 30*). This also reinforces our previous finding that the release of the Tc toxin YenTc from specialized *Y. entomophaga* cells occurs through the previously undescribed lytic action of the T10SS YenDF. Given the parallels observed, we propose to name the *S. marcescens* T10SS the *Serratia marcescens* death factor (SmaDF) for purposes of consistency.

## Discussion

Based on the entire body of our observations, we propose the following model for soldier cell differentiation and subsequent YenDF-mediated protein release. In a small percentage of isogenic cells at temperatures permissive for YenR expression, nutrient sensing, quorum sensing and further unknown factors cause an increase in the levels of YenR (**Fig. 4a**), possibly in the form of a positive autoregulatory loop as has been shown for bimodal expression of the *Bacillus subtilis* master spore regulator Spo0A (*35*). This enables YenR to drive transcription of soldier cell genes, resulting in the production of the YenDF components YeHln, YeEln, YeIspn and YeOspn, as well as a cytoplasmic cocktail of toxins including YenTc in a fully assembled holotoxin state (**Fig. 4b**). Upon an environmentally pH-triggered decrease of the PMF, small holin YeHln oligomers in the inner membrane either open pores sufficiently large to translocate the 15 kDa cytoplasmic endolysin YeEln into the periplasm (**Fig. 4c**) or flip YeEln across the membrane via a membrane weakening/flipping mechanism (*48*), where it cleaves peptidoglycan cross-links (*37*). Once YeEln has modified enough of the peptidoglycan sacculus that this no longer sterically hinders the spanins (*38*), the inner membrane anchored spanin YeIspn and the outer membrane anchored spanin YeOspn fold towards each other and ultimately fuse these membranes (**Fig. 4d**), which remain bound to the peptidoglycan layer by tethering Lpp proteins. The accompanying release of inner osmotic pressure (*49*) fires the cocktail of soldier cell toxins including YenTc into the external environment (**Fig. 4e**), where they can proceed to locate and act on their cellular targets.

**Figure 4.**
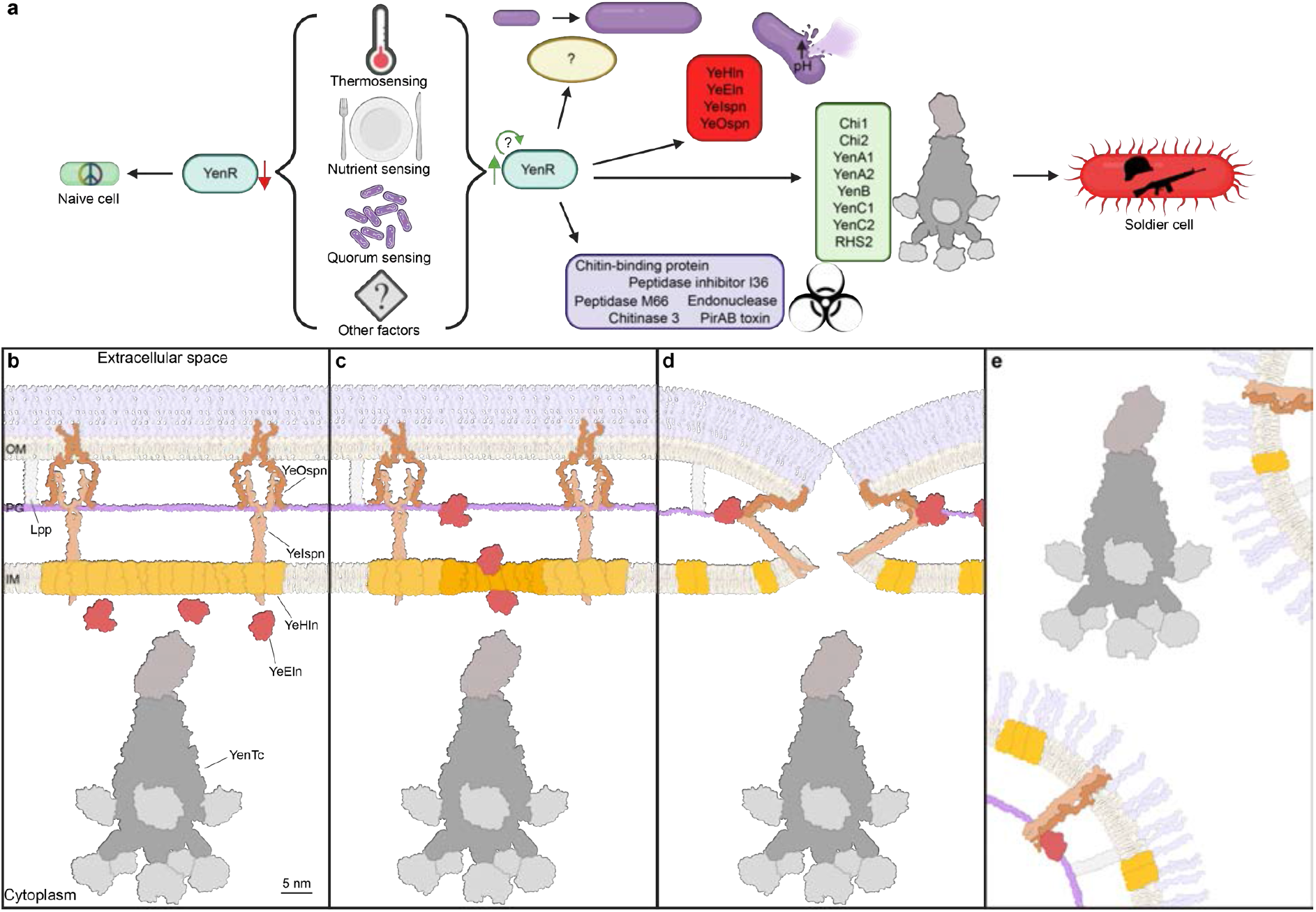
Model for soldier cell differentiation and subsequent YenDF-mediated YenTc release. **a**, At the lower temperatures that are conductive to YenR mRNA stability, synergy of several environmental cues causes an increase of YenR levels in a subset of cells, possibly aided by an autoregulatory positive feedback loop. This leads to differentiation into soldier cells by YenR-mediated transcription of YenTc components (green box), YenDF components (red box), additional toxins and virulence factors (purple box), and the characteristic enlarged phenotype of soldier cells by affecting an as yet undetermined set of genes. **b**, During YenR-mediated differentiation, soldier cells produce numerous toxins including YenTc, which is fully assembled in the cytoplasm. Components of the T10SS YenDF are produced simultaneously. YeEln remains in the cytoplasm, YeHln assembles into oligomeric rafts in the inner membrane, and the inner membrane embedded YeIspn forms a complex with the outer membrane embedded YeOspn. **c**, pH elevation that likely deteriorates the proton motive force either triggers the formation/opening of small pores in the YeHln rafts or enables pore-free YeHln mediated transmembrane flipping of YeEln, allowing YeEln to reach the periplasm and cleave peptidoglycan cross-links. **d**, Once enough cross-links have been cleaved, the spanin complex undergoes a conformational change that brings the inner and outer membranes into direct contact, leading to the fusion of the outer and inner membrane. **e**, The fused membranes collapse into loosely bound clusters of vesicles still tethered to peptidoglycan via the protein Lpp. The accompanying release of inner osmotic pressure propels YenTc and other soldier cell toxins into the surrounding environment.

In this study, we have discovered that the Gram-negative insect pathogen *Y. entomophaga* releases the Tc toxin YenTc and numerous other virulence factors into the surrounding environment using the type 10 secretion system YenDF. Furthermore, by also investigating the archetypal T10SS SmaDF from *S. marcescens* (**Figs. S11-S12**), we show that T10SSs mediate protein release by cell lysis rather than non-lytically as was previously hypothesized (*28, 29*). Due to its use of a holin/endolysin/spanin cassette, the mechanism described here has a degree of similarity to the lytic release of certain bacteriophage-encoded virulence factors such as Shiga toxins. The fundamental difference between the two is that in the latter case, production and release of such toxins are a byproduct of the phage infection cycle (and can therefore be activated by the SOS response pathway (*25*)), whereas here differentiated cells produce both the secretion system and its numerous cargoes in a dedicated, phage-independent manner that is correspondingly insensitive to the SOS response (**Fig. S5l**,**n**). T10SSs themselves however are likely to be evolutionary descendants of classical phage lysis cassettes that bacteria have adopted and repurposed for their own needs. Since the individual endolysin- and spanin-driven later stages of bacteriophage release in Gram-negative bacteria have so far never been visualized *in situ* using cryo-ET to the best of our knowledge, the tomographic data presented here are therefore of direct relevance to this deeply fundamental biological process.

Taking the abovementioned into account, this is the first example of anti-eukaryotic toxins using this type of very recently established secretion system (*28*), which is particularly apt for this case since the nearly ribosome-sized YenTc is fully assembled in the cytoplasm and would not fit through any of the previously described toxin secretion systems. These findings thereby establish a new avenue for extremely rapid release of toxins and other virulence factors previously not associated with any known secretion system, with virtually no upper or lower size limitation. Since they mediate the release of transcriptionally-coupled cargo proteins via lysis, T10SSs are not a secretion system in the classically used sense of the term. Nonetheless, they function as a secretion system on the populational level by allowing the entire bacterial population to benefit from the vast amounts of useful transcriptionally coupled cargo proteins synthesized by the small subset of T10SS-expressing cells. We therefore support the use of the term type 10 secretion system with respect to these specialized lysis cassettes.

Since cells pay the ultimate price for releasing proteins via YenDF, namely death, only a small specialized subset of what we term soldier cells employ it at any given time. How the decision to differentiate into a soldier cell is made has not yet been determined, although nutrient and quorum sensing play an important role and temperature sensing is crucial (**Figs. S2 and S5, Fig. 2f**). We discovered that the transcription factor YenR serves as the central temperature-sensitive switch of this bimodal differentiation, ensuring that the production of numerous toxins and their secretion system is synchronized (**Fig. 2**). The net result is a *Y. entomophaga* population that demonstrates traits such as differentiation and altruism reminiscent of eusocial systems, a far cry from the “bags of enzymes” that bacteria were once considered to be (*50*). From the perspective of costs and benefits to the entire bacterial population, it is rational that its few soldier cells are sacrificed to release as many host damaging factors as possible. This not only explains why soldier cells produce such a variety of virulence factors and toxins, but also why they tend to grow much larger than their non-differentiated isogenic brethren (**Fig. 1c, Fig. 2b and e**): this increases the total toxin carrying capacity of each specialized cell.

This also means that in a natural setting, production of YenTc and its secretion machinery is likely initiated only upon ingestion in response to the presence of host nutrients (**Fig. S5**), a situation mimicked by complex growth media. Since vomiting clears a significant portion of invading bacteria from the acidic anterior midgut (*34*), having a pH sensitive secretion mechanism helps prevent release of the toxic cocktail from soldier cells until they have reached the alkaline posterior midgut, their major theater of operations. At this point, chitinases and chitin-binding proteins of the cocktail would breach the chitinous peritrophic membrane (*51*) to enable access of YenTc and the other released toxins to the underlying midgut epithelial layer, leading to its complete dissolution and successful colonization of the insect (*34*) by the non-differentiated *Y. entomophaga* population.

*Y. entomophaga* serves as an excellent proxy for dangerous human pathogens that employ Tc toxins. Analysis of Tc toxin operons from e.g. *Yersinia pseudotuberculosis, Salmonella enterica* subsp. *houtenae, Yersinia enterocolitica*, and *Yersinia pestis* show that they also encode T10SSs similar to YenDF (**Fig. S13a-d**), indicating that these bacteria employ the same secretion mechanism as *Y. entomophaga*. Although the presence of a T10SS in *Y. enterocolitica* has been noticed before (*52*), its biological relevance has now been validated by demonstrating that without its T10SS this pathogen loses its ability to establish an infection in insects in a manner similar to a Tc toxin deletion mutant (*53*). Given that our reconstruction of the ancient *Y. pestis* Tc toxin operon from a Black Death victim (*54*) shows that it also encodes a T10SS (**Fig. S13d, inset**), this correspondingly raises questions to its role in the deadliest pandemic of human history (*55*).

The T10SS-mediated release of Tc toxins also fits well into the emerging paradigm of anti-eukaryotic toxin export by phage-derived proteins (*48*). Notable examples include release of the *Clostridium difficile* toxins A and B via a holin and endolysin (*56-58*), the holin-dependent export of large clostridial glucosylating toxins (*59, 60*) and the endolysin-dependent secretion of typhoid toxin (*61*). Notably, in all the cases described above, the current consensus is that the holin and/or endolysin-mediated export occurs via a non-lytic mechanism (*48, 56, 59-61*), although in light of the data presented here the potential use of suicidal soldier cell subpopulations also by these pathogens is a fascinating possibility that is worth investigating. Finally, combining the observation that YenTc deletion causes full loss of *Y. entomophaga* virulence (*22*) with our discovery that Tc toxins are produced by only few cells in a population leads to the startling insight that pathogen virulence can be determined by a small number of specialized soldier cells. If this is indeed found to be a more widespread phenomenon, then medical interventions that specifically target such specialized cells may represent a promising treatment strategy for many bacterial diseases.

## Materials and methods

### Bacterial strains and constructs

*Y. entomophaga* strains, *S. marcescens* strains and plasmids used in this study are provided in **Supplementary Table 1**.

### Cell growth and secretion assay conditions

*Y. entomophaga* type strain MH96 (*66*) and *S. marcescens* type strain BS 303 (also known as strain ATCC 13880) were obtained from the German Collection of Microorganisms and Cell Cultures GmbH (DSMZ). To test secretion or *Y. entomophaga* in different growth media, 10 mL of these were inoculated with 500 μL exponential phase culture grown in LB, then grown for 16 h at 16 °C. The media was separated from the cells by 5 min of centrifugation at 4000*g* and 0.22 μm filtration. Amicon concentrators (Merck) were used for concentrating the media to 250 μL. All samples were then normalized with 1x PBS to a volume equivalent to a cell OD_600_ of 4.0 and analyzed by stain-free SDS-PAGE. For secretion assays, 20 mL cells were grown in SOC growth medium for 16 h at 20 °C. Then the pH was elevated either by drop-wise addition of NaOH / a fixed volume of 1 M Tris-HCl pH 8.0, or resuspension of spun-down cells in 100 mM Tris-HCl pH 8.0, 100 mM NaCl buffer. *S. marcescens* cells were originally grown in either M9, LB or SOC media at 20 or 30 °C, with SOC at 20 or 30 °C used for later experiments. SOC media acidifies to the same extent in both *Y. entomophaga* and *S. marcescens* cultures. Ara-YenR and Ara-ChiR containing strains and the Ara-YeEln and Ara-ChiR plasmids were induced by addition of 0.5% L-arabinose during inoculation. For anaerobiosis-induced secretion, 50 mL induced and acidified cell cultures were left stationary after vigorously shaking during growth for up to 20 minutes in case of *Y. entomophaga* Ara-YenR and up to 60 minutes in case of *S. marcescens* Ara-ChiR.

### Targeted genomic editing of *Y. entomophaga* and *S. marcescens*

Cells were initially electroporated with an editing plasmid encoding λ-RED recombineering proteins, an I-SceI endonuclease, as well as a Cas9 protein plus a gRNA that has minimal off-target specificity in *Y. entomophaga* as determined by Cas-OFFinder (*67*). The cells were then transformed with a donor plasmid encoding an I-SceI cleavable fragment flanked by ∼300 bp homologous to the 5’ and 3’ sequences of the targeted area, containing an antibiotic resistance marker, a SacB counterselection marker, a 20 bp Cas9-gRNA targeting region, and a short repeat of the 5’ sequence. This setup allows modification of targeted genomic areas with optional subsequent Cas9-mediated excision of the antibiotic resistance marker (*68*), facilitating multiple sequential targeted genomic edits. The cells were then plated onto LB agar plates containing antibiotics for selection of the editing plasmid antibiotic resistance marker (kanamycin), the donor cassette antibiotic resistance marker (chloramphenicol) and 10% sucrose as a counterselection agent. Surviving colonies were then re-streaked onto an identical plate to eliminate background, and success of the genomic editing was validated by colony PCR and sequencing.

### Non-cryogenic cell imaging

For routine phase contrast and fluorescent imaging, 1 μL of *Y. entomophaga* or *S. marcescens* strain cultures were imaged on a glass slide at 20x using an EVOS M7000 microscope (Thermo Fisher Scientific). Confocal fluorescence microscopy timelapses of YenTc-sfGFP, as well as YenR-sfGFP and Ara-YenR YeHln-mCherry ΔYeEln/YeIspn/YeOspn cells before and after pH 8.0 induced secretion triggering was done on a glass slide at 40x using a LSM800 microscope (Zeiss) equipped with an Airyscan detector module (Zeiss).

### Fluorescence spectroscopy

For measuring the effect of different environmental conditions and quorum system knockouts on YenTc production, 3 biological replicates of YenTc-sfGFP cells with or without genomic knockouts of the acyl-homoserine-lactone synthase (for autoinducer-1 KO), the S-ribosylhomocysteine lyase (for autoinducer-2 KO), and the L-threonine 3-dehydrogenase (for autoinducer-3 KO) were grown in the conditions specified in **Fig. S5**. These were harvested, resuspended to OD_600_ 1.0 in 1x PBS, and used to measure the pre-secretion content of cellular YenTc-sfGFP. After 20 minutes incubation time, the cells were spun down and the supernatant was used to measure the content of secreted YenTc-sfGFP. Fluorescence emission spectra were recorded on a Spark spectrophotometer (Tecan) using an excitation wavelength of 470 nm and emission wavelength of 518 nm in a 2 × 2 read mode.

### Proteomic analysis using NanoHPLC-MS/MS

Biological triplicates of either the secreted protein fractions or the pre-secretion cytoplasmic contents from cells normalized to OD_600_ 4.0 were briefly run on a stain-free SDS-PAGE gel (Bio-Rad). After tryptic digestion and purification, the protein fragments were analyzed by nano-HPLC-MS/MS by using an Ultimate^™^ 3000 RSLC nano-HPLC system and a Hybrid-Orbitrap mass spectrometer (Q Exactive^™^ Plus) equipped with a nano-spray source (all from ThermoFisher Scientific). Briefly, the lyophilized tryptic peptides were suspended in 20 μL 0.1% TFA, and 3 μL of the samples were injected onto and enriched on a C18 PepMap 100 column (5 μm, 100 Å, 300 μm ID * 5 mm, Dionex, Germany) using 0.1% TFA, at a flow rate of 30 μL/min, for 5 min. Subsequently, the peptides were separated on a C18 PepMap 100 column (3 μm, 100 Å, 75 μm ID * 50 cm) using a linear gradient, starting with 95% solvent A/5% solvent B and increasing to 30.0% solvent B in 90 min using a flow rate of 300 nL/min followed by washing and re-equilibration of the column (solvent A: water containing 0.1% formic acid; solvent B: acetonitrile containing 0.1% formic acid). The nano-HPLC apparatus was coupled online with the mass spectrometer using a standard coated Pico Tip emitter (ID 20 μm, Tip-ID 10 μM, New Objective, Woburn, MA, USA). Signals in the mass range of m/z 300 to 1650 were acquired at a resolution of 70,000 for full scan, followed by up to ten high-energy collision-dissociation (HCD) MS/MS scans of the most intense at least doubly charged ions at a resolution of 17,500.

Relative protein quantification was performed by using MaxQuant (*69*) v.2.0.3.1, including the Andromeda search algorithm and searching the *Y. entomophaga* proteome of the UniProt database (downloaded Jan 2022). Briefly, an MS/MS ion search was performed for enzymatic trypsin cleavage, allowing two missed cleavages. Carbamidomethylation was set as a fixed protein modification, and oxidation of methionine and acetylation of the N-terminus were set as variable modifications. The mass accuracy was set to 20 ppm for the first search, and to 4.5 ppm for the second search. The false discovery rates for peptide and protein identification were set to 0.01. Only proteins for which at least two peptides were quantified were chosen for further validation. Relative quantification of proteins was performed by using the label-free quantification algorithm implemented in MaxQuant and the match-between-runs feature was activated.

Statistical data analysis of samples was performed using Perseus (*70*) v.1.6.14.0. Label-free quantification (LFQ) intensities were log-transformed (log2); replicate samples were grouped together. Proteins had to be quantified at least three times in at least one of the groups of a comparison to be retained for further analysis. Missing values were imputed using small normal distributed values (width 0.3, down shift 1.8 for the datasets involving the *Y. entomophaga* Ara-YenR strainand *S. marcescens* strains; width 0.3, down shift 2.0 for the dataset assessing the *Y. entomophaga* secreted vs non-secreted protein fractions) and a two-sided *t*-test (significance threshold: -log2 fold change > 1.5 for all datasets; p-value < 0.02 for the datasets involving the *Y. entomophaga* Ara-YenR strain; p-value < 0.05 for the dataset assessing the *Y. entomophaga* secreted vs non-secreted protein fractions; p-value < 0.01 for the datasets assessing the *S. marcescens* strains) was performed. Proteins that were statistically significant outliers were considered as hits. Volcano plots were generated using VolcaNoseR (*71*).

UniProt accession numbers (parenthesized) of statistically significant hits from *Y. entomophaga* according to the above criteria, which are of major biological importance to this study are as follows: YenA1 (B6A877), YenA2 (B6A878), Chi1 (B6A876), Chi2 (B6A879), YenB (B6A880), YenC1 (B6A881), YenC2 (B6A882), RHS2 (A0A2D0TC51), Cpb (A0A3S6EXR6), PirA (A0A3S6F007), PirB (A0A3S6F043), PiI36 (A0A3S6F569), NucA (A0A3S6F4M5), Chi3 (A0A3S6F1Q8), StcE (A0A3S6EYX4), Tlh (A0A3S6F052), YeEln (A0A3S6F4L4), YenIspn (A0A3S6F4Q6), YenR (A0A3S6F5G2). Such hits from *S. marcescens* are: ChiR (M4SHQ2), ChiX (A0A349ZDQ1), ChiY (A0A379YYR5), ChiA (A0A379Y6D9), ChiB (P11797), ChiD (A0A380ANW3), Cbp21 (O83009).

### RT-qPCR detection

For quantifying the effect of temperature on YenR transcript levels, 3 biological replicates of either *Y. entomophaga* Ara-YenR or YenR-sfGFP strain cells were grown at 16 °C or 37 °C in SOC medium overnight with shaking, for the Ara-YenR strain in presence of 0.5% L-arabinose. A volume of culture corresponding to a final OD_600_ of 0.1 was added to 750 μL TRIzol reagent (Thermo Fisher Scientific) per sample, and total RNA was purified according to manufacturer protocols. The RNA was then directly treated with the DNA-free DNAse treatment kit (Thermo Fisher Scientific) according to manufacturer protocols in order to remove any contaminating DNA from the total RNA preparations. RT-qPCR was carried out immediately afterwards in a CFX96 system (Bio-Rad) using a Power SYBR Green RNA-to-C_T_ 1-Step Kit (Thermo Fisher Scientific), which also contains RNase inhibitor and ROX dye for passive referencing. 100 nM primers with sequences CCCTCGCAAAGATTGTAATTCA and CACTGGTTAATCATGCGTCAA were used to target YenR. 100 nM primers with sequences CCTTACATACTTCCAAACACCC and CCAAAACTGACTATCTGATGCG were used to target YmoA in the same samples, which served as a temperature-invariant control. Raw data was analyzed using the CFX Manager software (Bio-Rad), with a threshold value of 605 RFU used to determine C_q_. The acquired data were then assessed for significance using paired t-tests and further processed using Prism 9 (GraphPad).

### Bioinformatic analyses

Identification of secretion signal sequences for YenR-controlled toxins and virulence factors was performed using SignalP 6.0 (*72*). The sequence logo for the final 30 bp before the start codon of YenR-controlled genes (as per **Fig. 2d**) was calculated using WebLogo 3.7.12 (*73*) following multiple sequence alignment via Clustal Omega (*74*) with combined iterations and maximum guide tree / HMM iterations set to 5. The promoter region of the polycistronic YenDF structural component operon was also included in this analysis. For reconstruction of the YpeTc operon of ancient *Y. pestis* from a Black Death victim, genomic reads from Illumina run SRR341961 of the original study (*54*) were assembled into contigs using Ray Meta (*75*), which were in turn assembled into a scaffold using CSAR-Web (*76*) with the YpeTc operon of *Y. pestis* strain KIM10+ as a reference.

### Bacterial vitrification

Overnight cultures of *Y. entomophaga* strains were spun down for 4 min at 4000*g* and resuspended to an OD_600_ of 20 in 1x PBS buffer containing pre-washed 10 nm BSA-NanoGold tracer (Aurion). In case of the Ara-YenR strain, 10 μL was directly applied to glow-discharged Quantifoil R1/4 Au-SiO_2_ 200 grids and incubated for 10 minutes in a Vitrobot Mark IV plunger (Thermo Fisher Scientific) set to 100% humidity and 22 °C prior to blotting and plunge-freezing. 3 μL of the other three strains were applied to identically treated grids in identical plunger conditions following 30 min incubation in the 1x PBS-NanoGold buffer. After a waiting time of 60 s, all grids were blotted from both sides for 32 s using blot force 5. After 0.5 s drain time, the grids were vitrified by plunge-freezing into liquid ethane. For *Y. entomophaga* Ara-YenR ΔYenEln/YeIspn/YeOspn cells intended for whole cell tomography, the OD_600_ was adjusted to 5 and blot time to 10 s, and Quantifoil R2/1 Au-SiO_2_ 200 grids were used. Overnight cultures of the *S. marcescens* Ara-ChiR strain at an OD_600_ of 8.5 were transferred to a non-shaking Eppendorf tube for 1 h, 10 nm BSA-NanoGold tracer (Aurion) was added, and the sample was then applied to glow-discharged Quantifoil R1/4 Au-SiO_2_ 200 grids. Blotting conditions were identical to those used for *Y. entomophaga*. Grids containing *Y. entomophaga* Ara-YenR strain cells and *S. marcescens* Ara-ChiR strain cells were clipped with standard AutoGrids (Thermo Fisher Scientific) and directly used for further data acquisition as the cells were thin enough without additional cryo-FIB milling.

### Lamella preparation by cryo-FIB milling

Grids containing *Y. entomophaga* Ara-YenR ΔYenDF, Ara-YenR ΔYenEln/YeIspn/YeOspn, or Ara-YenR ΔYeIspn/YeOspn strain cells were clipped in cryo-FIB-specific AutoGrids (Thermo Fisher Scientific) with alignment markers and a cut-out for milling at shallow angles. Clipped grids were transferred to an Aquilos 2 cryo-FIB/SEM dual beam microscope (Thermo Fisher Scientific). Lamella preparation was performed as previously described (*77*). Briefly, after platinum sputter coating and deposition of metalloorganic platinum, clusters of bacteria cells were targeted for a four-step milling procedure using decreasing ion beam currents from 0.5 nA to 50 pA. Milling angles of 6-10° relative to the grid were used. Lamellae were milled to a thickness range of 50-100 nm.

### Cryo-ET data acquisition

Once ready for imaging, all grids were transferred into a Titan Krios transmission electron microscope (Thermo Fisher Scientific) operated at 300 kV, equipped with a K3 camera and a BioQuantum energy filter (Gatan). Images were acquired with SerialEM (*78*). Overview images were acquired at 6500x nominal magnification to identify regions for cryo-ET data acquisition at higher magnification. Images used as references for batch data acquisition were also acquired at this magnification. Tilt series were acquired at 64,000x (pixel size 1.48Å) for milled *Y. entomophaga* Ara-YenR ΔYenDF, Ara-YenR ΔYenEln/YeIspn/YeOspn, and Ara-YenR ΔYeIspn/YeOspn cells and at 42,000x (pixel size 2.32 Å) for *Y. entomophaga* Ara-YenR and *S. marcescens* Ara-ChiR cells using a script based on a dose-symmetric tilt scheme (*79*) at defoci ranging from -5 to -8 μm. The stage was tilted from -60° to +60° relative to the lamella plane at 3° increments. Each tilt series was exposed to a total dose of 120-140 e^-^/Å^2^. Tilt series for intact *Y. entomophaga* Ara-YenR ΔYenEln/YeIspn/YeOspn cells were acquired at 33,000x magnification (pixel size 2.861 Å) with a Volta phase plate (*80*). Before acquiring each tilt series, a new phase plate position was activated for 2 minutes. A dose-symmetric tilt scheme from -54° to +54° with a 3° increment was used for data acquisition. A defocus of -0.5 μm was applied. Each tilt series was exposed to a total dose of 77 e^-^/Å^2^.

### Tomogram reconstruction and sub-tomogram averaging of YenTc

Acquired movie frames were motion-corrected and combined into stacks of tilt series using Warp (*81*). The stacks were aligned and reconstructed using IMOD (*82*). During alignment, patch tracking was used in tilt series of milled cells and fiducial-marker tracking was used in tilt series of non-milled cells. The tomograms were 4x binned and low-pass filtered to 60 Å or 100 Å for better visualization using EMAN2 (*83*). Alternatively, denoising by an implementation of cryo-CARE (*84*) was used for the same purpose. To obtain a structure of YenTc, 167 particles were manually picked from 12 tomograms of Ara-YenR cells. The sub-tomograms were extracted from 4x binned tomograms with a box size of 100 pixels (928 Å) using RELION 3.0(*85*). The sub-tomograms were aligned to a spherical reference and averaged over iterations with C5 symmetry in RELION (**Fig. S10**).

Making use of the attractive properties of Ara-YenR ΔYeIspn/YeOspn cell spheroplasts, intracellular YenTc holotoxin particles were located in these cells using TomoTwin (*65*) (**Fig. S9**) and used for structure generation via subtomogram averaging. Particles were automatically picked with a pre-release of TomoTwin 0.3 (*65*) using a clustering workflow. First, all 7 tomograms were rescaled to 15 Å to make the YenTc particles fit into the TomoTwin static box-size of 37×37×37. Next, a single tomogram was embedded with the latest general model (version 052022) which resulted in 4015980 embedding vectors. To remove embedding vectors corresponding to background volumes, the median embedding vector was calculated and all embeddings that had a cosine similarity with the median embedding vector higher than 0 were discarded. From the remaining 527451 embedding vectors a 2D UMAP was calculated (**Fig. S9**), with the highlighted cluster corresponding to the YenTc particles. The average of all embedding vectors belonging to this cluster gave the reference embedding that was used further picking. The 6 remaining tomograms were then embedded with TomoTwin and the reference was the used to locate the YenTc particles. Using a confidence threshold of 0.846 gave a total number of 528 particles.The sub-tomograms were then extracted from 4x binned tomograms with a box size of 128 pixels (760 Å) using RELION 3.0. The sub-tomograms were aligned to a spherical reference and averaged over iterations with C5 symmetry in RELION (**Fig. S10**).

### Tomogram annotation and figure production

In order to visualize the morphological changes accompanying YenDF component action as well as the position and assembly status of YenTc, densities of YenTc, the inner membrane, peptidoglycan layer, outer membrane and fused membranes (in AraYenR tomograms) as well as exemplary cell envelope-spanning proteins (in AraYenR tomograms) were annotated on a slice-to-slice basis in reconstructed tomograms using Dragonfly (Object Research Systems). Weaker densities were masked during this process, which culminated in generation of 3D segmented volumes.

ChimeraX (*86*) was used for **Fig. S10c-f. Fig. 4a** and **Fig. S1b** were created using Biorender.com.

## Acknowledgements

We thank K. Vogel-Bachmayr and S. Bergbrede for wet lab technical support, O. Hofnagel and D. Prumbaum for electron microscopy technical support, and A. Brockmeyer and W. Hecker for mass spectrometric technical support. Special thanks go to P. Njenga Ng‘Ang‘A, W. Oosterheert and G. Rice for fruitful discussions. This study was supported by funding from the Max Planck Society (to S.R.).

## Author contributions

O.S. and S.R. initiated and designed the project. O.S. carried out most of the experiments. Z.W., O.S. and T.W. acquired and analyzed tomographic data. P.J. analyzed mass spectrometric data. L.K. and O.S. generated bacterial strains. S.R. supervised the project. O.S. and S.R. wrote the manuscript with input from all co-authors.

## Data availability statement

The raw data generated during the current study are available from the corresponding authors upon reasonable request. Source data (includes unprocessed SDS-PAGE gels, mass spectrometry proteomics data, and raw data for graphs) are provided with this paper in the Source Data file. The raw mass spectrometry proteomics data have been deposited to the ProteomeXchange Consortium via the MassIVE partner repository with the dataset identifiers MSV00089961 / PXD035561 (secreted fraction vs. non-secreted fraction of *Y. entomophaga* cultures), MSV000089964 / PXD035573 (induced Ara-YenR vs. WT or vs. non-induced Ara-YenR *Y. entomophaga*), and MSV000091191, PXD039813 (secreted fraction / pre-secretion fraction of induced *S. marcescens* Ara-ChiR vs. *S. marcescens* WT or vs. induced ΔSmaDF *S. marcescens*). YenTc cryo-ET structures from the post-endolysin and post-spanin states have been deposited in the Electron Microscopy Data Bank under accession numbers EMD-16618 and EMD-15403, respectively. Representative tomograms for *Y. entomophaga* are deposited under accession numbers EMD-15404 (pre-secretion state), EMD-15405 (post-holin state, FIB-milled), EMD-16619 (post-holin state, intact cells), EMD-15406 (post-endolysin state) and EMD-15407 (post-spanin state). Representative tomograms for *S. marcescens* in the post-spanin state are deposited under accession numbers EMD-16538 and EMD-16539.

