## Supplementary Figures and Tables for "Specialized pathogenic cells release Tc toxins using a type 10 secretion system"

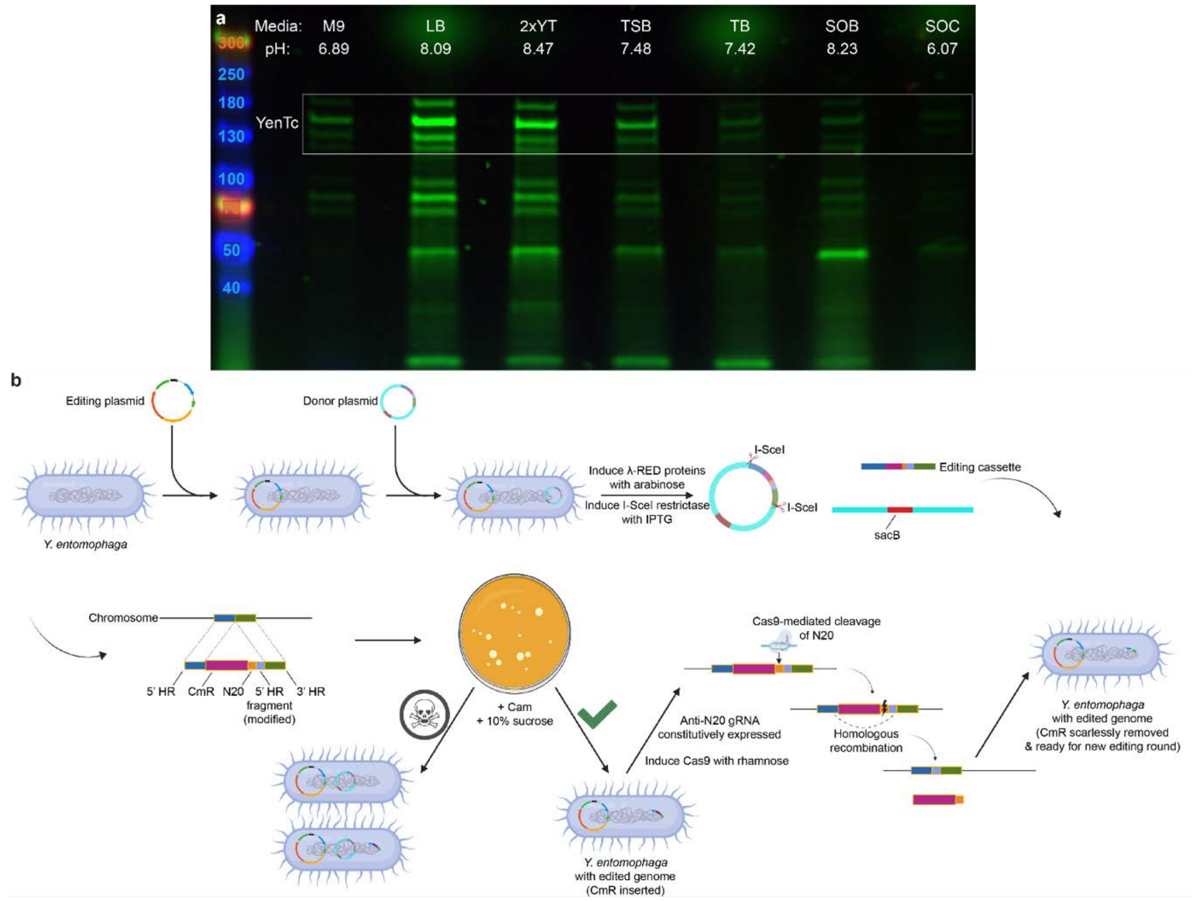


**Supplementary Figure 1 | Fundamentals of the controlled setup used to find and characterize the YenTc secretion system. a,** Secreted protein fractions from *Y. entomophaga* cultured in various growth media. Secretion is inhibited in media that acidifies during cell growth. Bands corresponding to YenA1, YenA2 and YenB are boxed for clarity. **b,** Schematic representation of the targeted scarless genomic editing protocol established for *Y. entomophaga*. The same protocol was also used to modify the genome of *S. marcescens*.

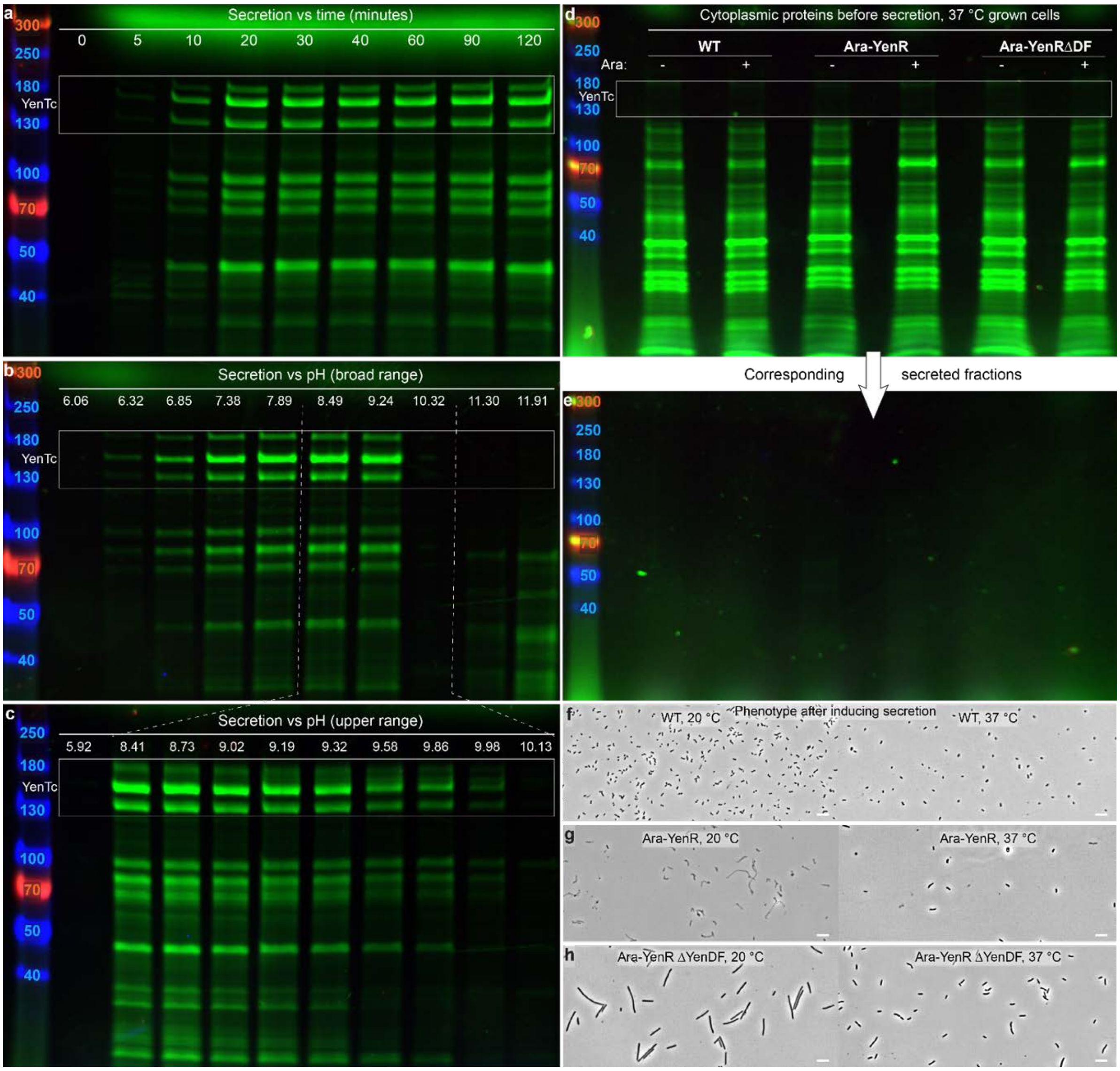


**Supplementary Figure 2 | Soldier cells secrete YenTc within a few minutes and at near-neutral to slightly alkaline pH, only at lower temperatures. a,** *Y. entomophaga* cells grown in acidified SOC media rapidly secrete when pH is adjusted to 8.0. Bands corresponding to YenA1, YenA2 and YenB are boxed for clarity. **b,** Broad range screening of the pH dependence of secretion, assessed by adjusting the pH of acidified SOC media to the values indicated. Non-specific cell lysis seen in the two most alkaline samples demonstrates the upper limit of *Y. entomophaga* pH tolerance. **c,** Narrow range screening of the pH dependence of secretion to determine the upper pH limit of soldier cell specific secretion. **d-e,** *Y. entomophaga* does not produce or secrete toxins when grown at 37 °C as opposed to the usual 20 °C, even when YenR is induced. **f-g**, The phenotype of WT and Ara-YenR strain cells grown at 20 °C and 37 °C after pH-induced secretion.

22

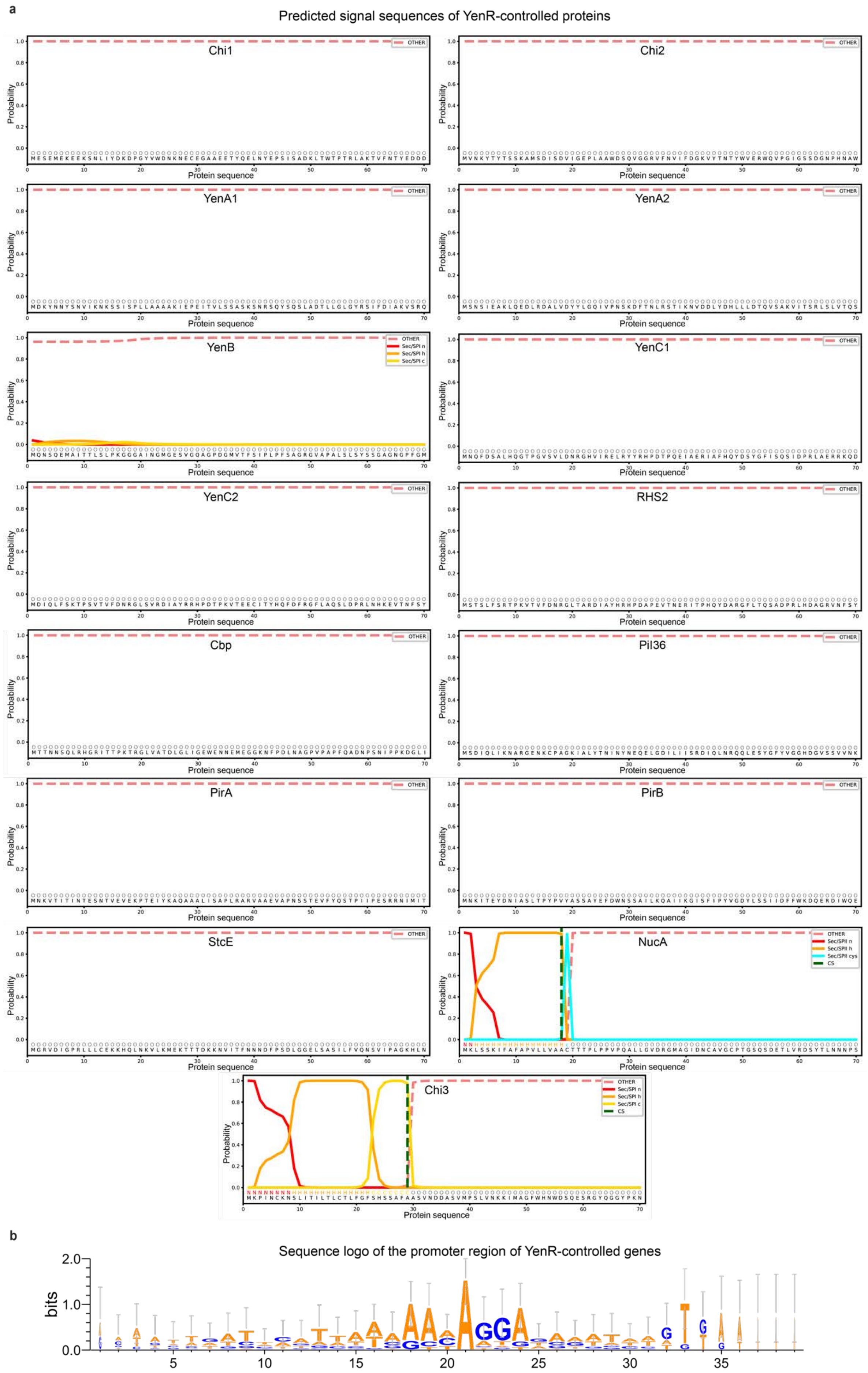
a
Predicted signal sequences of YenR-controlled proteins
Chi1
Chi2
YenA1
YenA2
YenB
YenC1
YenC2
RHS2
Cbp
Pil36
PirA
PirB
StcE
NucA
Chi3
OTHER
Sec/SPI n
Sec/SPI h
Sec/SPI c
Sec/SPII cys
CS
Probability
Protein sequence
b
Sequence logo of the promoter region of YenR-controlled genes
bits

**Supplementary Figure 3 | Nearly all YenR-controlled toxins and virulence factors lack an established signal sequence and have a putative YenR promoter recognition sequence. a,** Predictions of secretion signal sequences for YenR-controlled toxins and virulence factors show that nearly all are unable to utilize alternative export pathways due to lack of a secretion signal sequence. Of those that do, the nuclease NucA contains disulfide bridges that require an oxidative environment for maturation, and periplasmically localized chitinases (as likely the case for Chi3) have been suggested to target soluble oligosaccharides that enter the cell through porins (*62*). The relevant UniProt accession numbers are provided in the Methods section. **b,** Sequence logo derived from the promoter region of YenR-controlled toxins, virulence factors and the YenDF structural component operon indicates an overrepresented sequence that may function as a YenR binding site.

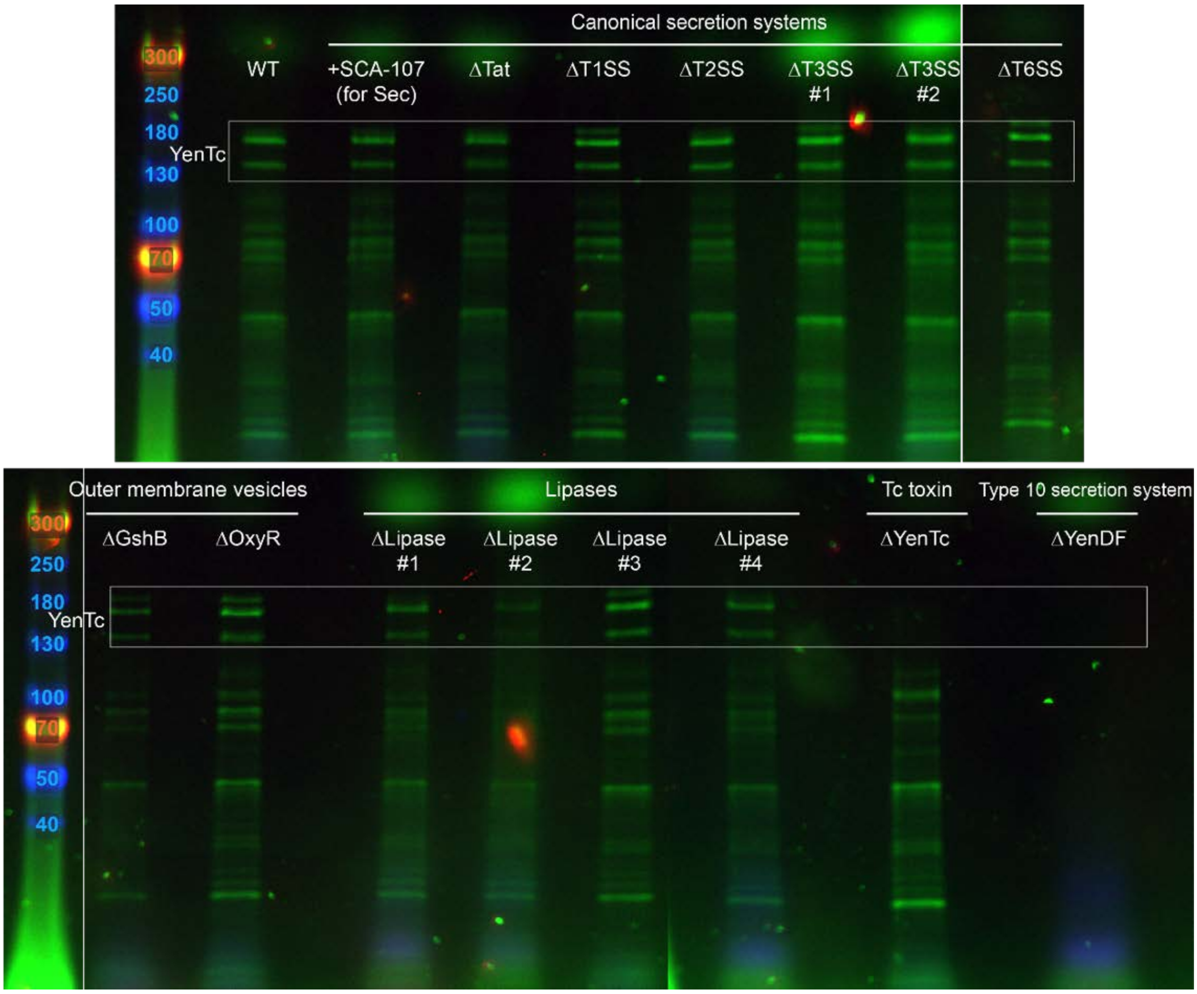


**Supplementary Figure 4 | A T10SS was identified as the major pathway for *Y. entomophaga* protein secretion.** *Y. entomophaga* knockout strains of established anti-eukaryotic toxin secretion pathways or previously proposed Tc toxin export pathways have WT-like secretion levels, while the knockout of YenDF, a novel T10SS, abolished protein export completely. The Sec export pathway is essential and non-removable and was therefore blocked by the specific chemical inhibitor SCA-107 (*63*). While no single gene controlling OMV formation is known, absence of OxyR and GshB very strongly decreased *E. coli* OMV formation levels (*64*). *Y. entomophaga* does not encode a direct Pdl1 lipase homologue that was proposed to be a *Photorhabdus* Tc toxin release factor, so four proteins with lipase-like domains that might serve as a proxy were targeted, as was YenTc itself to see if it has autotransporter-like capabilities. Bands corresponding to YenA1, YenA2 and YenB are boxed for clarity.

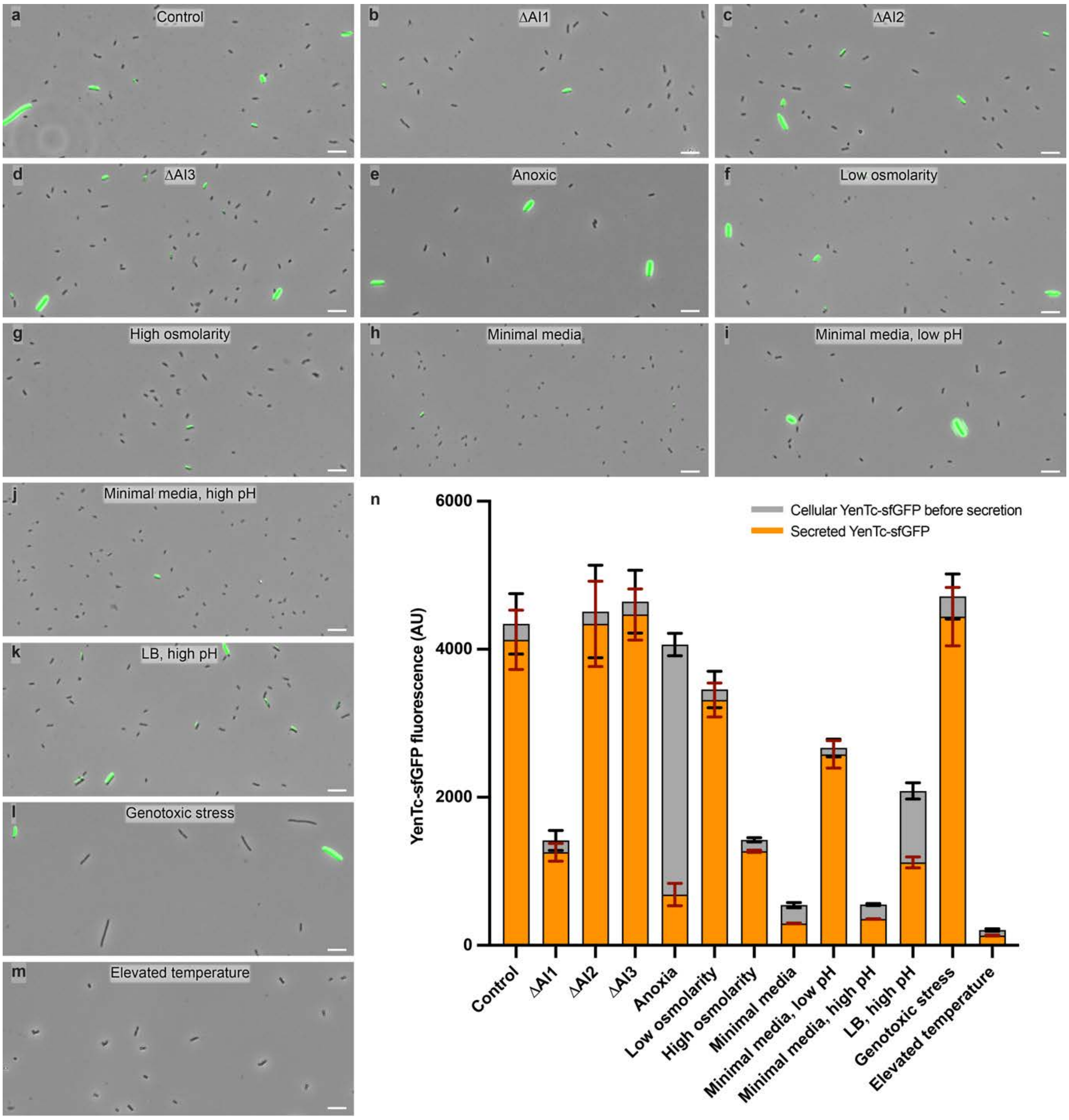


**Supplementary Figure 5 | Differentiation into soldier cells is primarily controlled by thermosensing, nutrient availability, and autoinducer-1 quorum sensing molecules. a,** A genomic YenTc-sfGFP fusion was used as a qualitative marker of differentiation into soldier cells (green). *Y. entomophaga* was grown in the acidifying growth media SOC at 20 °C unless noted otherwise. **b-d,** Knockout of the acyl-homoserine-lactone synthase (UniProt accession number: A0A3S6EWZ2), the S-ribosylhomocysteine lyase (UniProt accession number: A0A3S6F351), and the L-threonine 3-dehydrogenase (UniProt accession number: A0A3S6EZE1) to test for the essentiality of autoinducer-1, -2 and -3 type quorum sensing molecules in differentiation, respectively. Absence of autoinducer-1 molecules decreases the number of soldier cells noticeably. **e,** Growth in a pure nitrogen atmosphere to test the effect of anoxia on differentiation. Interestingly, anoxically grown cells demonstrate defective YenTc secretion, but unimpaired YenTc production (**n**). **f-g,** Growth in SOC media prepared with 0 or 570 mM NaCl to test the effect of decreased and increased osmolarity on differentiation. Normal SOC media contains 190 mM NaCl. **h,** Growth in M9 minimal media to test for the essentiality of complex organic molecules for differentiation. In the absence of complex organic molecules, the cells tend to adopt minimal dimensions and strongly reduce differentiation into soldier cells. **i-j,** Same as in (**e**) but pre-adjusted to pH 6.10 / pH 7.70 to respectively test for the influence of acidic and alkaline pH on differentiation in absence of complex organic compounds. Acidic, but not alkaline pH, stimulates production of YenTc (**n**). **k,** Growth in LB media, which increases its pH value to 7.50 after cell growth, to test how alkaline pH in presence of complex organic compounds affects differentiation compared to (**a**) and (**j**). **l,** Growth in media supplemented with 200 ng/mL mitomycin C to test the effect of genotoxin stress on differentiation and the relevance of the SOS response pathway, which tends to generate enlarged cells by inhibition of cell division. **m,** Growth at 37 °C to test the effect of elevated temperature on differentiation. Given the lack of observed YenTc-sfGFP expression, of all factors tested here high temperature affects the propensity of naive cells to differentiate the most. Scale bars: 10 µm. **n,** Differentiation into soldier cells (grey bars) and their secretory ability (superimposed orange bars) in conditions from (**a-m**) was quantified by measuring fluorescence of YenTc-sfGFP in the pre-secretion and secreted fractions, respectively, of density-normalized cells. Data is shown as mean ± standard deviation, n = 3 biological replicates.

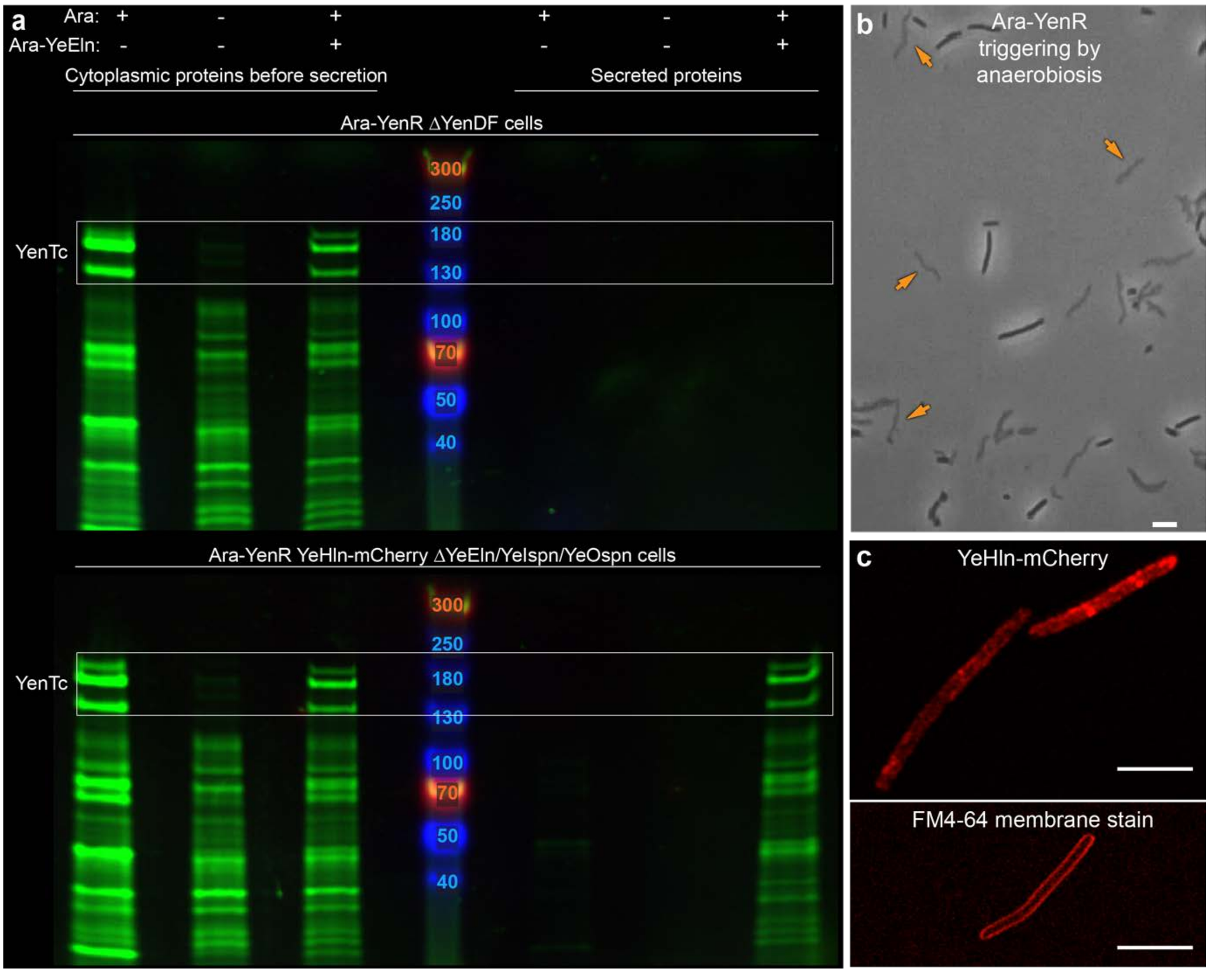


**Supplementary Figure 6 | pH change triggers YeHln, which concentrates into numerous small foci.** **a,** YeHln-mCherry enables protein release in response to elevation of pH in Ara-YenR YeHln-mCherry ΔYeEln/YeIspn/YeOspn cells. In this particular strain, Ara-YeEln was produced from a plasmid in lieu of its (deleted) genomic copy to avoid interference of the fused mCherry tag to YeEln expression, whereas in the remaining experiments of this study YeEln is expressed via its native gene. For similar reasons, YeIspn and YeOspn were also deleted and given the toxicity we observed for plasmid-borne spanin overexpression, shearing forces applied to the bacteria during sample preparation were used as a replacement for their action, as has been previously documented (*38*). Bands corresponding to YenA1, YenA2 and YenB are boxed for clarity. **b,** Aerobic shaking cultures of Ara-YenR cells can be stimulated to undergo YenDF-mediated secretion by immobility-induced anaerobiosis. Orange arrowheads denote examples of cells that already underwent YenDF-mediated secretion. See Fig. 2b for an intact reference. Scale bar: 5 µm. **c**, Confocal fluorescence microscopy of Ara-YenR YeHln-mCherry ΔYeEln/YeIspn/YeOspn cells shows that small YeHln raft-like oligomers are abundantly distributed across the bacterial membrane which contrasts to the few large lesions observed for triggered phage holin S105 (*43*), and the smooth membrane signal produced by FM4-64 staining of AraYenR ΔYenDF cells. The YeHln-mCherry fusion presented here was shown to be fully functional by the experiments depicted in (**a**). Scale bars: 5 µm.

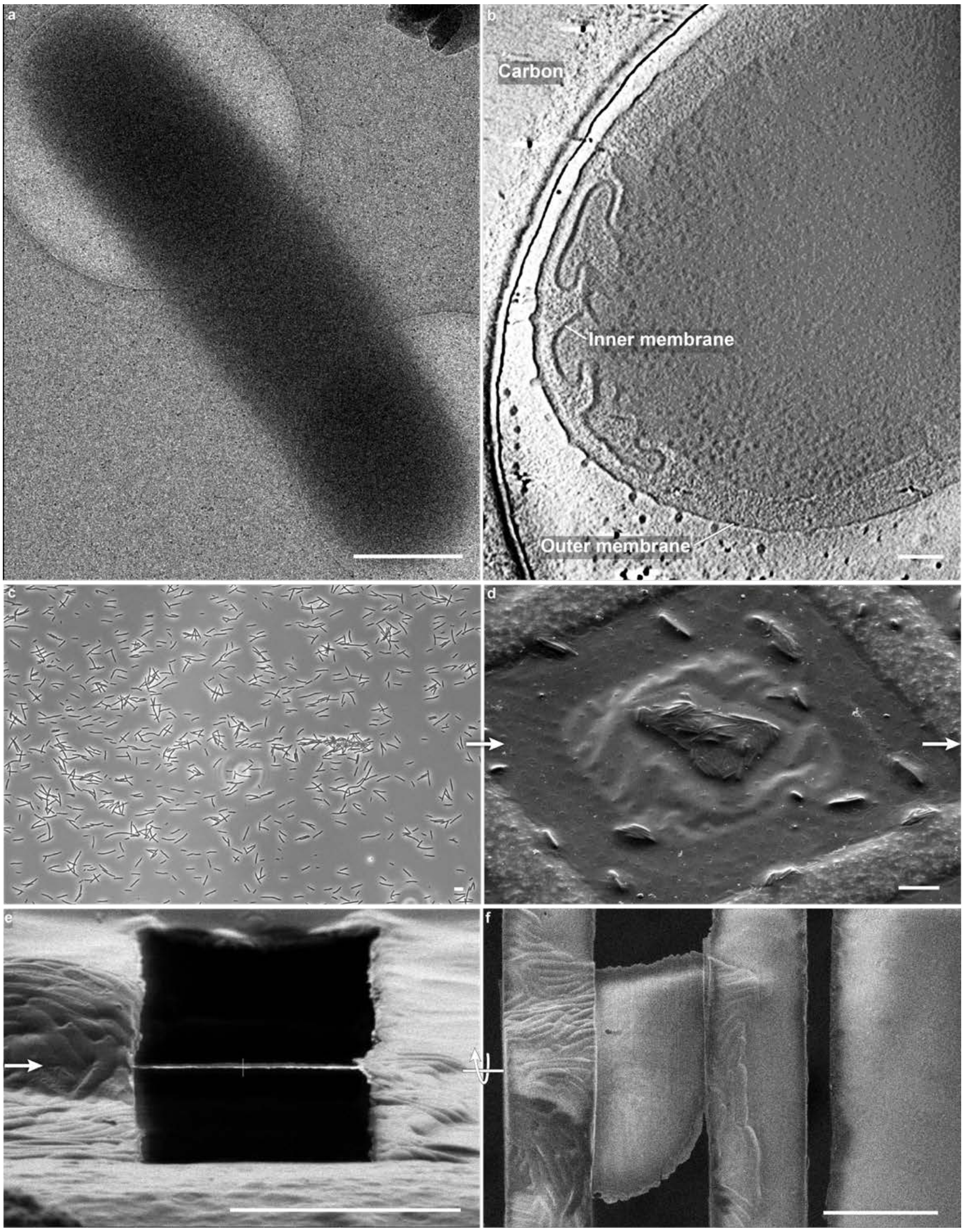


**Supplementary Figure 7 | The importance and process of bacterial sample preparation by cryo-focused ion beam milling.** **a**, A TEM overview image of an intact, unmilled, pH-induced Ara-YenR ΔYeEln/YeIspn/YeOspn cell, showing poor contrast due to the large size of *Y. entomophaga* (soldier) cells. Scale bar: 1 µm. **b**, A tomogram obtained from an intact pH-induced Ara-YenR ΔYeEln/YeIspn/YeOspn cell showing numerous minor inner membrane invaginations attributed to YeHln action. Notice how even with use of a Volta phase plate, only limited features are observable compared to the tomograms acquired on FIB-milled samples shown in other figures. Scale bar: 100 nm. **c**, Induced Ara-YenR ΔYenDF cells used to investigate the pre-secretion state of soldier cells, prior to vitrification. Scale bar: 10 µm. **d**, SEM micrograph of plunge-frozen, pH-triggered Ara-YenR ΔYenDF cells on an EM grid, prior to lamella generation. Scale bar: 10 µm. **e-f**, SEM micrograph of an Ara-YenR ΔYenDF cell lamella milled with a focused ion beam, side and top view. Scale bars: 10 µm.

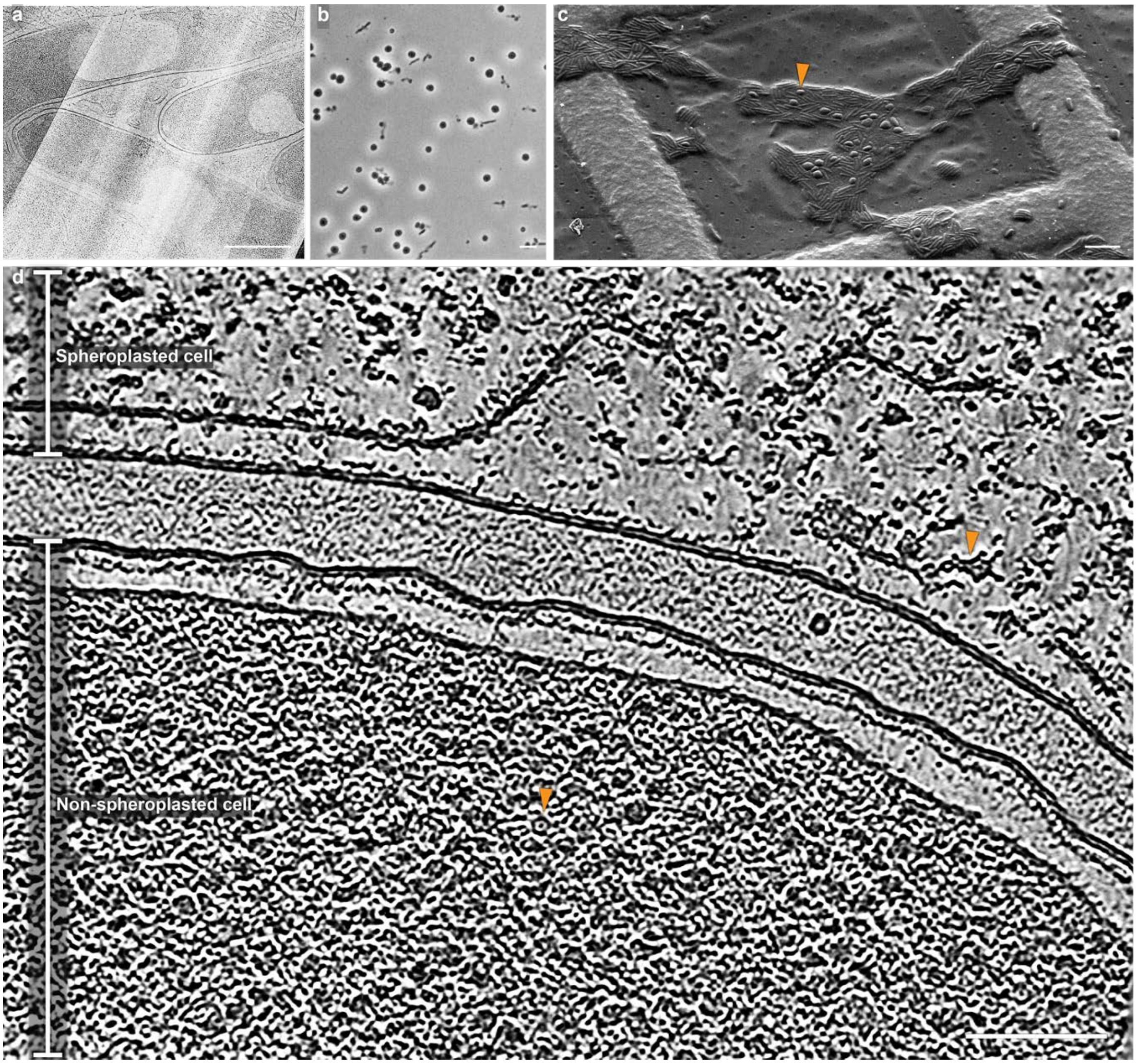


**Supplementary Figure 8 | Spanin knockout soldier cells form spheroplasts with attractive qualities for *in situ* tomographic protein analysis. a,** During a ~<30 min window after pH-induced activation, enzymatic activity of YeHln-translocated YeEln causes Ara-YenR ΔYeIspn/YeOspn strain cells to exhibit severe inward bending of their inner membranes, as seen in this overview image of a cryo-FIB milled lamella. Scale bar: 1 µm **b,** Eventually such cells transform into spheroplasts. Scale bar: 10 µm. **c,** SEM micrograph of plunge-frozen AraYenR ΔYeIspn/YeOspn cells that were pH-triggered on an EM grid, prior to lamella generation. Arrowhead: an example spheroplast. Scale bar: 10 µm. **d,** A single slice from a AraYenR ΔYeIspn/YeOspn cell tomogram illustrating the difference in interior protein density of a spheroplasted cell compared to a non-spheroplasted cell. Notice the ultrastructural changes to the spheroplast inner membrane that occur as a result of cell expansion. Arrowheads: an example YenTc holotoxin from a spheroplast (side view) and non-spheroplast (top view). Scale bar: 100 nm.

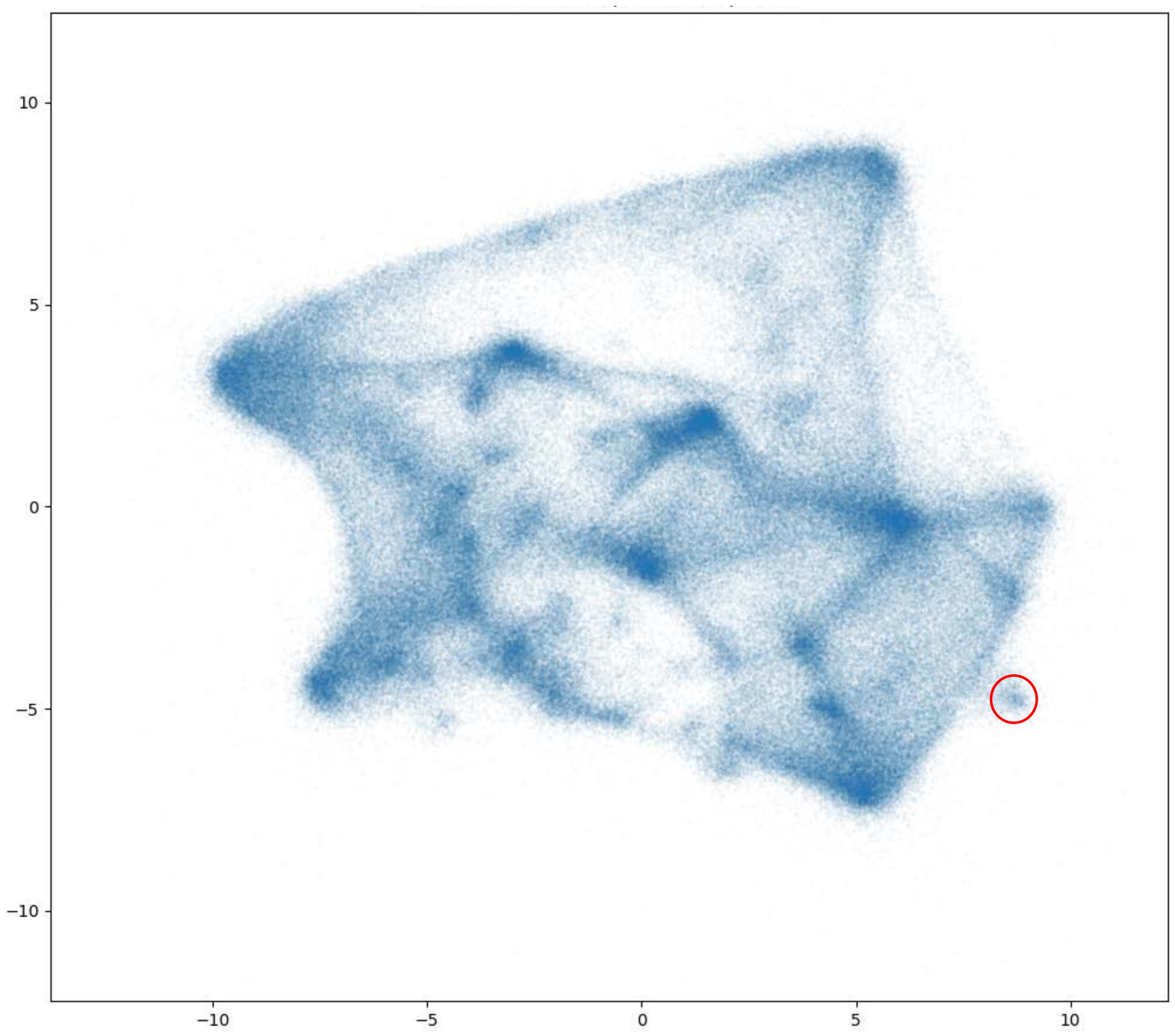


**Supplementary Figure 9 | The YenTc cluster from a single embedded tomogram as seen in a TomoTwin UMAP.** The UMAP was calculated on the median filtered embeddings, and average of all embedding vectors from this cluster was to create a reference embedding used for unsupervised picking of YenTc in the remaining tomograms.

121

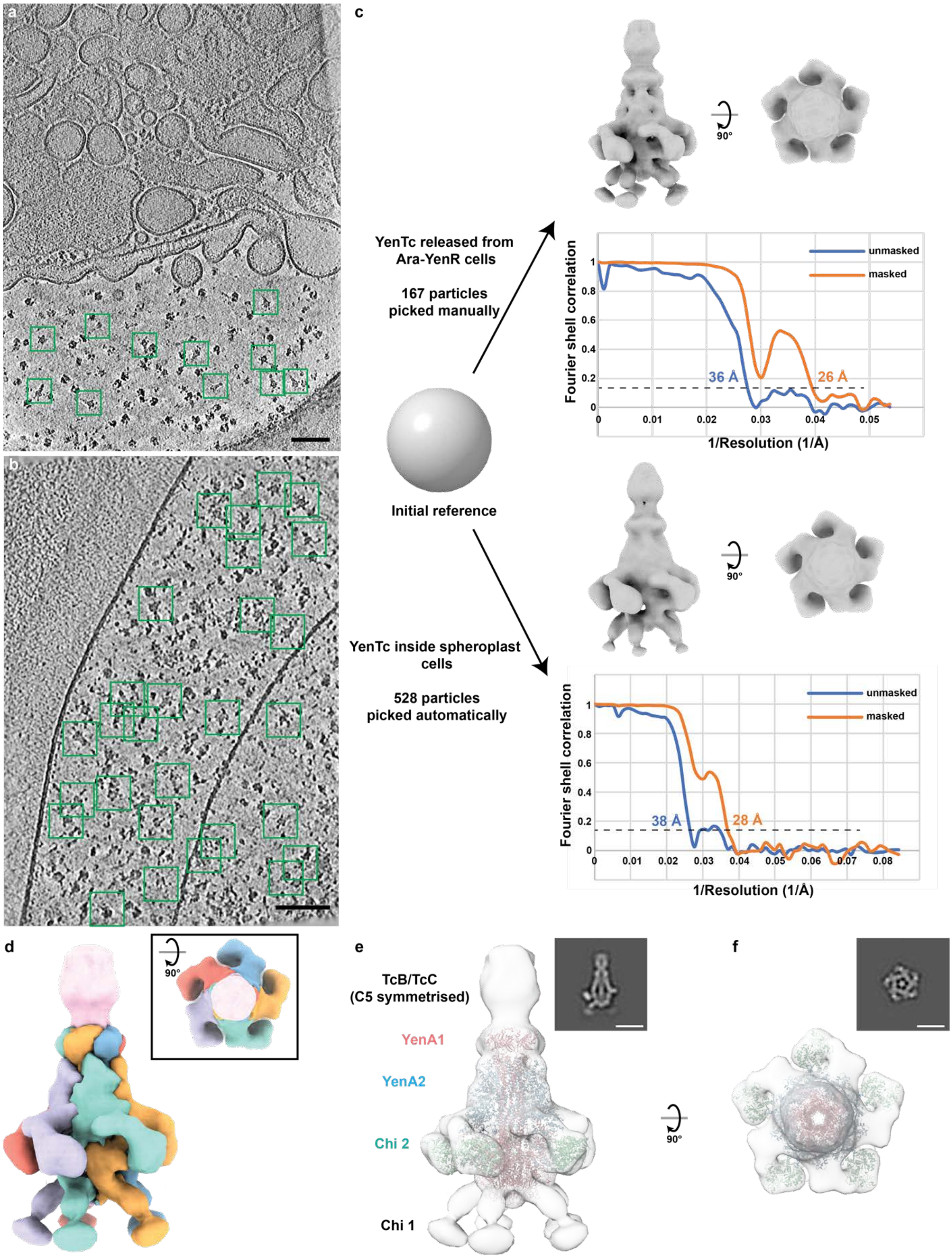
a
b
c
YenTc released from Ara-YenR cells
167 particles picked manually
Initial reference
YenTc inside spheroplast cells
528 particles picked automatically
90°
unmasked
masked
Fourier shell correlation
1/Resolution (1/Å)
36 Å
26 Å
38 Å
28 Å
d
e
f
TcB/TcC (C5 symmetrised)
YenA1
YenA2
Chi 2
Chi 1

**Supplementary Figure 10 | Subtomogram averaging of YenTc before and after release by soldier cells. a,** A 9 nm thick slice of a representative tomogram of Ara-YenR cells. YenTc particles manually picked for subtomogram averaging are highlighted by green boxes. Scale bar: 100 nm. **b,** A 6-nm thick slice of a representative tomogram of Ara-YenR ΔYeIspn/YeOspn spheroplast cells. YenTc particles automatically picked by TomoTwin (*65*) are highlighted by green boxes. Scale bar: 100 nm. **c,** Subtomogram averaging of YenTc released from Ara-YenR cells (top) and localized intracellularly in Ara-YenR ΔYeIspn/YeOspn cells (bottom), with corresponding gold-standard FSC curves of the YenTc structures. The dip in the FSC curve at 33 Å corresponds to the first zero of the CTF curve at a defocus of -5.5 µm, which is the defoci of the majority of the particles. **d,** The structure of YenTc from Ara-YenR cells with individually coloured protomers of TcA. The TcB/TcC components are coloured in pink. Inset: top view. **e-f**, Side and top view of the YenTc structure fitted with a structural model of the YenTc TcA component (PDB: 6OGD) (*10*). Insets are cross-sections of the structure. Scale bar: 20 nm.

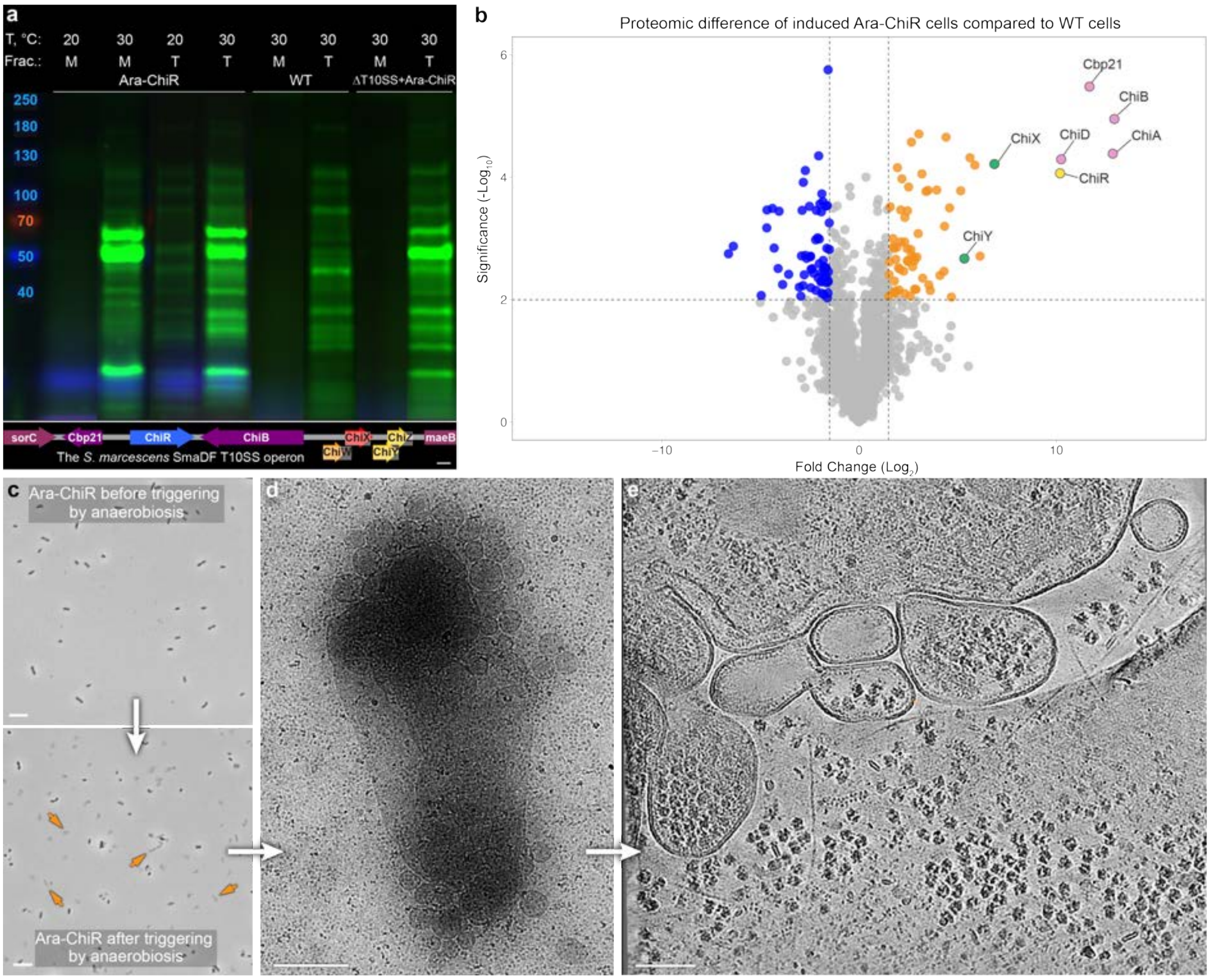


**Supplementary Figure 11 | The *Serratia marcescens* T10SS exports proteins via a lytic mode of action. a**, Arabinose induction of the Ara-ChiR strain leads to the temperature-sensitive production and secretion of chitinolytic proteins compared to wild type (WT) cells, whilst removing the SmaDF T10SS abolishes their secretion. Since the WT variant of the *S. marcescens* type strain does not express the ChiR T10SS/chitinolytic machinery regulator under the laboratory conditions tested, to switch on its production we either replaced the native ChiR promotor with arabinose-inducible machinery (in case of the Ara-ChiR strain) or transformed cells with an arabinose-inducible Ara-ChiR plasmid (in case of the SmaDF KO strain), similarly to what was earlier done by use of an arabinose-inducible plasmid (*30*). Fractions: M, secreted into growth media, T, total intracellular protein. Inset: structure of the SmaDF operon. Like classical phage lysis cassettes and the *Y. entomophaga* YenDF T10SS presented here, the *S. marcescens* SmaDF T10SS is composed of a holin (ChiW), endolysin (ChiX), i-spanin (ChiY) and o-spanin (ChiZ). Bimodal expression (*29*) of the T10SS from *S. marcescens* strain Db10 was shown to be controlled by the nearby LysR-type transcription factor ChiR (*30*), which also regulates the levels of chitinolytic proteins Cbp21, ChiA, ChiB and ChiC. The T10SS components of the type strain BS 303 used here have 95-99% sequence identity to those of the publicly unavailable strain Db10 investigated previously (*29*). Scale bar: 200 bp. **b**, Volcano plot showing significant enrichment (t-test p-value < 0.01) of chitinolytic and T10SS components in the total protein fraction of induced Ara-ChiR cells compared to ChiR-nonexpressing WT cells. This shows that ChiR not only controls the expression of both the T10SS and several chitinolytic machinery components as previously reported (*29, 30*), but also reveals the previously undescribed control of chitinase ChiD expression by ChiR. Chitinases ChiA and ChiD are encoded genomically distally from the T10SS, similarly to most *Y. entomophaga* YenR-controlled genes. Pink: chitinolytic components. Green: structural components of the SmaDF T10SS. Orange: enriched intracellular proteins. Yellow: the regulatory protein ChiR. The relevant UniProt accession numbers are provided in the Methods section. n = 3 biological replicates each. The full proteomic datasets used to generate these figures are available in the Source Data file. **c**, Aerobic shaking cultures of Ara-ChiR cells can be stimulated to undergo SmaDF-mediated secretion by immobility-induced anaerobiosis. Orange arrowheads denote examples of cells that already underwent SmaDF-mediated secretion. Scale bars: 10 µm. **d**, A TEM overview image of an Ara-ChiR cell after SmaDF-mediated secretion. Scale bar: 500 nm. **e**, Tomogram slice of an Ara-ChiR cell after secretion, corresponding to a post-spanin action state. Note the similarity to the phenotype observed for *Y. entomophaga* after YenDF spanin action in Fig. 3d, including formation of unimembrane vesicle clusters and release of cytoplasmic proteins into the environment. The observed SmaDF-mediated lysis explains the abundance of cytoplasmic and membrane proteins previously observed in the supernatant of *S. marcescens* strain Db10 cells, which was at the time attributed to sensitivity of the mass spectrometry technique (*29*). Scale bar: 100 nm.

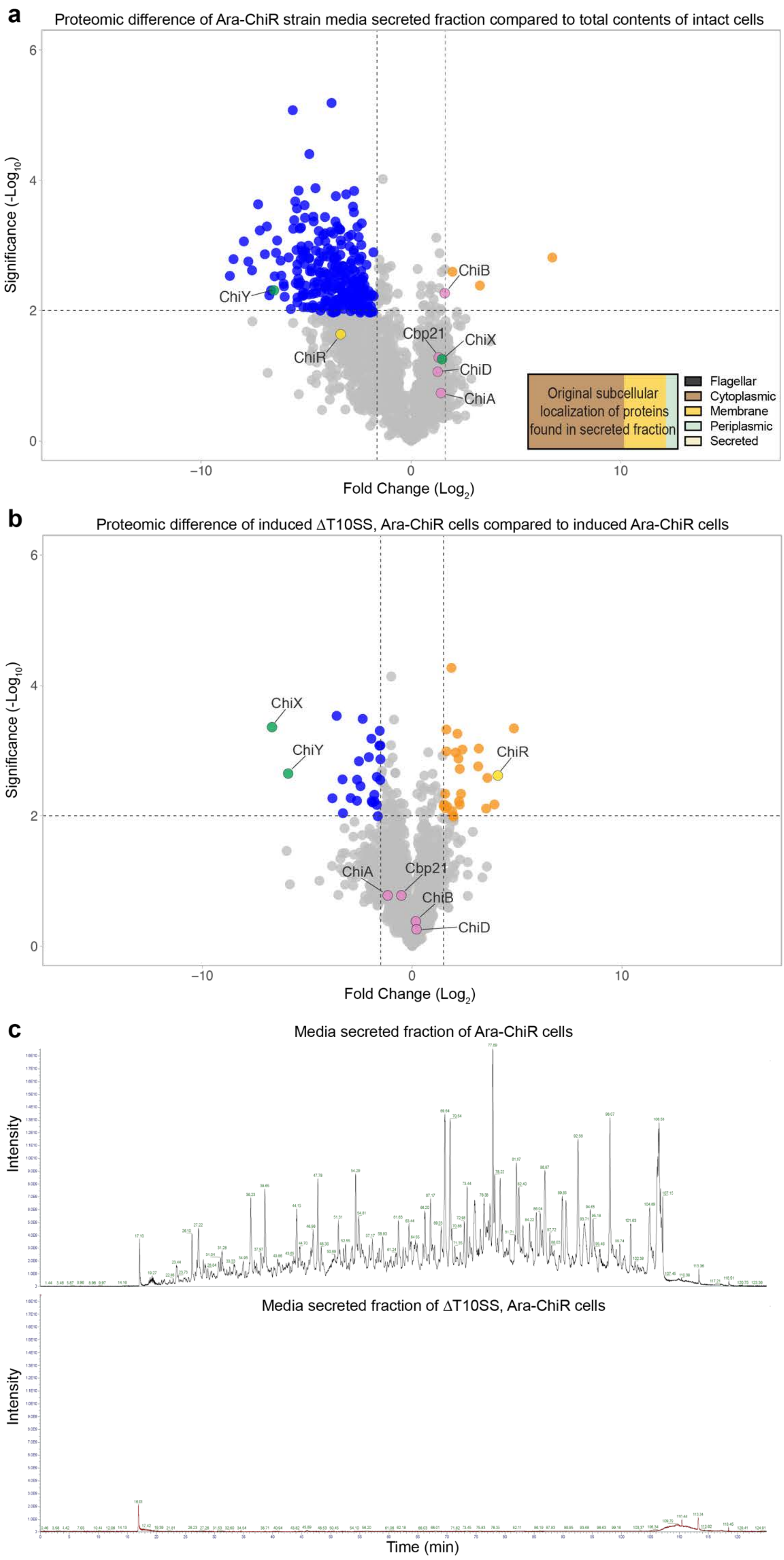
a
Proteomic difference of Ara-ChiR strain media secreted fraction compared to total contents of intact cells
Significance (-Log10)
Fold Change (Log2)
ChiY
ChiB
Cbp21
ChiX
ChiR
ChiD
ChiA
Original subcellular localization of proteins found in secreted fraction
Flagellar
Cytoplasmic
Membrane
Periplasmic
Secreted
b
Proteomic difference of induced ΔT10SS, Ara-ChiR cells compared to induced Ara-ChiR cells
Significance (-Log10)
Fold Change (Log2)
ChiX
ChiY
ChiR
ChiA
Cbp21
ChiB
ChiD
c
Media secreted fraction of Ara-ChiR cells
Intensity
Media secreted fraction of ΔT10SS, Ara-ChiR cells
Intensity
Time (min)

**Supplementary Figure 12 | The *Serratia marcescens* T10SS enables release of most intracellular proteins into the surrounding environment. a-b,** Volcano plot of induced Ara-ChiR cells showing the (a) overall similarity of the protein fraction already secreted into growth media compared to the total protein fraction of still intact cells (inset: original subcellular localizations of those proteins found in the secreted fraction for which such information is available (443 of 1817 total hits), and (b) overall similarity of total cell proteomes from Ara-YenR cells compared to SmaDF T10SS knockout cells expressing an arabinose-inducible version of ChiR from a plasmid. Comparison of the secreted fraction to the pre-secretion fraction shows that they are highly similar (grey circles correspond to most hits), as are the cytoplasmic fractions of the Ara-ChiR cells compared to the SmaDF knockout cells expressing a plasmid-encoded ChiR, which is produced in the latter at higher levels than the genomically encoded version in the former. A t-test p-value < 0.01 was used, and colors match Fig. S11b. n = 3 biological replicates each. The full proteomic datasets used to generate these figures are available in the Source Data file. **c,** Total ion chromatograms from the media secreted fractions of Ara-ChiR cells and SmaDF T10SS knockout cells expressing an arabinose-inducible version of ChiR from a plasmid, showing the abundance of proteins released into the media via action of SmaDF and their complete absence therefrom upon deletion of this T10SS.

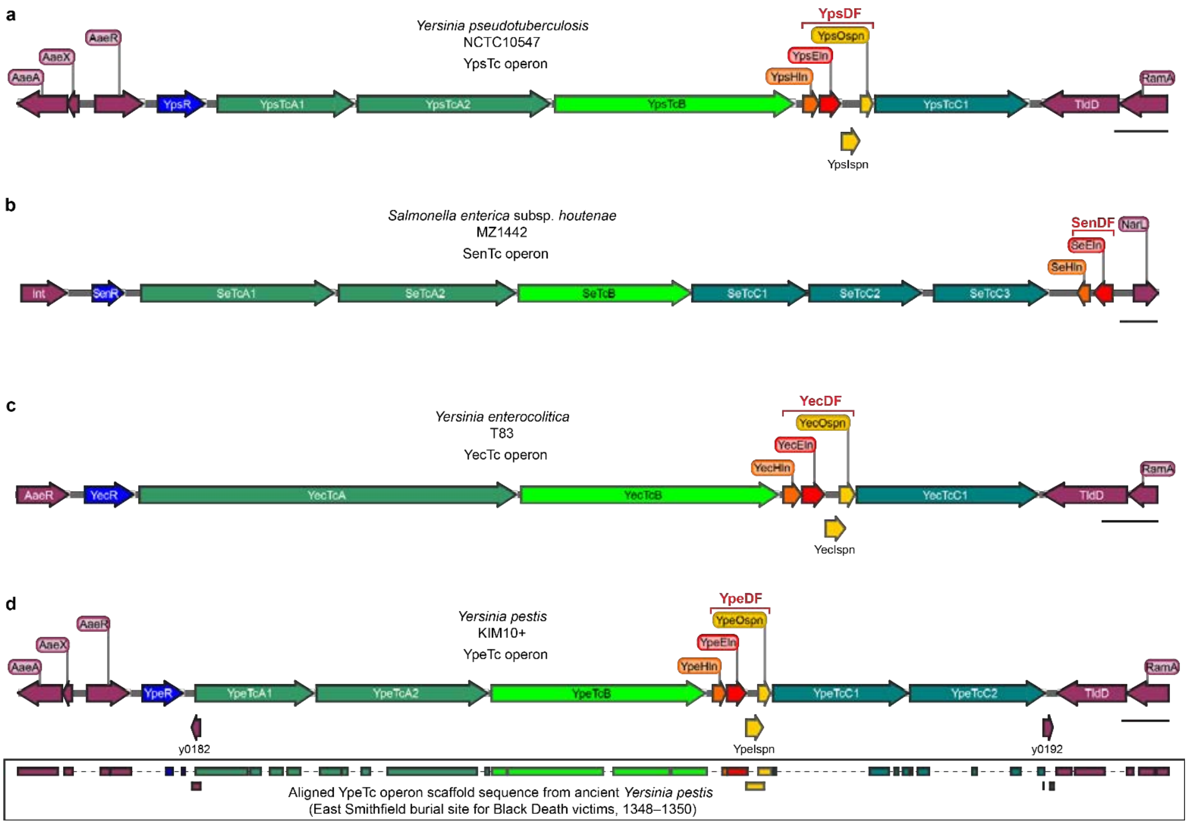


**Supplementary Figure 13 | Major human pathogens have type 10 secretion systems associated with Tc toxin genomic regions. a-d,** The Tc toxin genomic regions from *Yersinia pseudotuberculosis* (strain NCTC10547), *Salmonella enterica* subsp. *houtenae* (strain MZ1442), *Yersinia enterocolitica* (strain T83) and *Yersinia pestis* (strain KIM10+), with the embedded T10SSs designated as YpsDF, SenDF, YecDF and YpeDF in analogy to YenDF. The holin component is shown in orange, endolysin in red, spanins in yellow, putative transcriptional regulator of the Tc toxin operon in blue, TcA components in dark green, TcB in light green, TcC in blue-green, and genes not related to Tc toxins in burgundy. Inset of (**d**): the aligned YpeTc operon scaffold sequence from a London victim of the Black Death pandemic, ca. 1348-1350 (*54*). Regions shown with a dashed line had no available contigs due to degradation of ancient DNA. Scale bars: 1000 bp.

**Movie 1 | Synchronous YenDF-mediated controlled lysis of pH-induced *Y. entomophaga* Ara-YenR cells.**

**Movie 2 | Annotated tomogram overview of a *Y. entomophaga* pre-secretion state cell.**

**Movie 3 | Annotated tomogram overview of a *Y. entomophaga* post-holin action cell.**

**Movie 4 | Annotated tomogram overview of a *Y. entomophaga* post-endolysin action cell.**

**Movie 5 | Annotated tomogram overview of a *Y. entomophaga* post-spanin action cell.**

**Movie 6 | SmaDF-mediated controlled lysis of *S. marcescens* induced by anaerobiosis.**

**Movie 7 | Non-annotated tomogram overview of a *S. marcescens* post-spanin action cell (1).**

**Movie 8 | Non-annotated tomogram overview of a *S. marcescens* post-spanin action cell (2).**

**SI Table 1.** *Y. entomophaga* strains and plasmids used in this study.

**a.** *Y. entomophaga* strains used in this study.

| ***Y. entomophaga* strain** | **Description** | **Source** |
|---|---|---|
| MH96 | Wild type strain | German Collection of Microorganisms and Cell Cultures GmbH (DSMZ) |
| YenA1-sfGFP | sfGFP fusion after residue 37 of the YenA1 subunit of YenTc | This work |
| YenR-sfGFP | sfGFP fusion at the C-terminus of YenR | This work |
| ΔYeHln | YeHln to CmR | This work |
| ΔYeEln | YeEln to CmR | This work |
| ΔYeIspn/YeOspn | YeIspn and YeOspn to CmR | This work |
| ΔYenDF | YeHln, YeEln, YeIspn and YeOspn to CmR | This work |
| Ara-YenR | araC and AraBAD promoter inserted directly before YenR | This work |
| Ara-YenR ΔYenDF | Ara-YenR and ΔYenDF combination strain | This work |
| Ara-YenR ΔYeHln | Ara-YenR and ΔYeHln combination strain | This work |
| Ara-YenR ΔYeEln | Ara-YenR and ΔYeEln combination strain | This work |
| Ara-YenR ΔYeIspn/YeOspn | Ara-YenR and ΔYeIspn/YeOspn combination strain | This work |
| Ara-YenR YeHln-mCherry ΔYeEln/YeIspn/YeOspn | Ara-YenR combination strain with YeHln an mCherry fusion after a 30 aa linker, plus ΔYeEln/YeIspn/YeOspn | This work |
| Ara-YenR YenA1-sfGFP | Ara-YenR and YenA1-sfGFP combination strain | This work |
| ΔTat | TatA-TatD to CmR | This work |
| ΔT1SS | TolC to CmR | This work |
| ΔT2SS | PL78_RS08960-PL78_RS08990 to CmR | This work |
| ΔT3SS #1 | PL78_RS19995-PL78_RS1969 to CmR | This work |
| ΔT3SS #2 | PL78_RS18105-PL78_RS18250 to CmR | This work |
| ΔT6SS | PL78_RS00905-PL78_RS19390 to CmR | This work |
| ΔGshB | GshB to CmR | This work |
| ΔOxyR | OxyR to CmR | This work |
| ΔLipase #1 | PL78_RS09630 to CmR | This work |
| ΔLipase #2 | PL78_RS18445 to CmR | This work |
| ΔLipase #3 | PL78_RS18020 to CmR | This work |
| ΔLipase #4 | PL78_RS00835 to CmR | This work |
| ΔYenTc | Chi1-YenC2 to CmR | This work |

| | | |
|---|---|---|
| YenA1-sfGFP ΔAI1 | YenA1-sfGFP and acyl-homoserine-lactone synthase to GmR combination strain | This work |
| YenA1-sfGFP ΔAI2 | YenA1-sfGFP and S-ribosylhomocysteine lyase to AmpR combination strain | This work |
| YenA1-sfGFP ΔAI3 | YenA1-sfGFP and L-threonine 3-dehydrogenase to TcR combination strain | This work |

*S. marcescens* strains used in this study.

| ***S. marcescens*** **strain** | **Description** | **Source** |
|---|---|---|
| BS 303 (aka ATCC 13880) | Wild type strain | German Collection of Microorganisms and Cell Cultures GmbH (DSMZ) |
| Ara-ChiR | araC and AraBAD promoter inserted directly before ChiR | This work |
| ΔSmaDF | ChiW, ChiX, ChiY and ChiZ to CmR | This work |

**b.** Plasmids used in this study.

| **Plasmid** | **Description** | **Source** |
|---|---|---|
| Ara-YeEln | Arabinose-inducible YeEln | This work |
| Ara-ChiR | Arabinose-inducible ChiR | This work |
| pMultiEdit-v4 | Helper plasmid encoding arabinose-inducible λ-RED proteins, IPTG-inducible I-SceI restrictase, rhamnose-inducible Cas9 and constitutively expressed anti-N20 gRNA. The N20 is a sequence that has minimal off-target specificity in *Y. entomophaga*. | This work |
| pDonor | SacB-containing donor backbone plasmid for targeted genomic editing of regions of interest. | This work |
